# Delivery of CAR-T Cells in a Transient Injectable Stimulatory Hydrogel Niche Improves Treatment of Solid Tumors

**DOI:** 10.1101/2021.12.20.473538

**Authors:** Abigail K. Grosskopf, Louai Labanieh, Dorota D. Klysz, Gillie A. Roth, Peng Xu, Omokolade Adebowale, Emily C. Gale, Carolyn K. Jons, John H. Klich, Jerry Yan, Caitlin L. Maikawa, Santiago Correa, Ben S. Ou, Andrea I. d’Aquino, Jennifer R. Cochran, Ovijit Chaudhuri, Crystal L. Mackall, Eric A. Appel

**Affiliations:** Department of Chemical Engineering, Stanford University, Stanford, CA, 94305, USA; Department of Bioengineering, Stanford University, Stanford, CA, 94305, USA; Center for Cancer Cell Therapy, Stanford Cancer Institute, Stanford University School of Medicine, Stanford, CA, 94305, USA; Department of Biochemistry, Stanford University, Stanford, CA, 94305, USA; Department of Materials Science and Engineering, Stanford University, Stanford, CA, 94305, USA; Department of Mechanical Engineering Engineering, Stanford University, Stanford, CA, 94305, USA; Department of Pediatrics, Stanford University School of Medicine, Stanford, CA, 94305, USA; Stanford Cancer Institute, Stanford University School of Medicine, Stanford, CA, 94305, US; Department of Medicine, Stanford University School of Medicine, Stanford, CA, 94305, USA; ChEM-H Institute, Stanford University, Stanford, CA, 94305, USA; Department of Pediatrics - Endocrinology, Stanford University, Stanford, CA, 94305, USA

## Abstract

Adoptive cell therapy (ACT) has proven to be highly effective in treating blood cancers such as B cell malignancies, but traditional approaches to ACT are poorly effective in treating the multifarious solid tumors observed clinically. Locoregional cell delivery methods have shown promising results in treating solid tumors compared to standard intravenous delivery methods, but the approaches that have been described to date have several critical drawbacks ranging from complex manufacturing and poor modularity to challenging adminstration. In this work, we develop a simple-to-implement self-assembled and injectable hydrogel material for the controlled co-delivery of CAR-T cells and stimulatory cytokines that improves treatment of solid tumors. We evaluate a range of hydrogel formulations to optimize the creation of a transient inflammatory niche that affords sustained exposure of CAR-T cells and cytokines. This facile approach yields increased CAR-T cell expansion, induces a more tumor-reactive CAR-T phenotype, and improves efficacy in treating solid tumors in mice.

## Introduction

Adoptive cell therapy (ACT) is a promising new strategy to treat cancer which has been shown to be highly efficacious in the treatment of blood cancers. In the ACT process, immune cells are collected from a patient, isolated and engineered with receptors to recognize and eradicate cancer, expanded and then infused back into the patient for treatment. Chimeric Antigen Receptor (CAR) T cells are engineered to target an antigen that is over-expressed on cancer cells. This strategy has been widely effective in treating blood cancers including B-cell leukemias and lymphomas, but has seen limited success in treating the multifarious solid tumors observed clinically^1^. Several therapies for treating B-cell malignancies have recently been approved by the US Food & Drug Administration, and numerous concerted efforts are underway to translate this success to solid tumor applications^2^.

Currently, CAR-T cells are primarily delivered through intravenous (IV) infusion, which is effective in treating blood cancers because the T cells do not need to find and penetrate a solid tumor microenviroment. Unfortunately, T cells administered in this way often become trapped in the lungs and exhibit poor infiltration into solid tumors. Indeed, large numbers of CAR-T cells are typically required for current treatment strategies to be successful, which requires costly, complex, and labor intensive *ex vivo* expansion of the CAR-T cells that can inhibit their effector potential when transferred for treatment^3, 4^. In contrast, locoregional cell delivery methods – particularly those exploiting biomaterial scaffolds – can improve upon systemic delivery for treatment of solid tumors and elicit increased local expansion of T cells at the tumor site, improved tumor infiltration and enhanced efficacy^5–8^. Unfortunately, the biomaterials scaffolds developed for locoregional ACT to date often require invasive surgical implantation procedures to reach tumor sites, hindering translation, and typically limiting the use of these approaches to tumor resection or post-surgery applications^6, 8, 9^. Furthermore, many of the biomaterials reported to date do not degrade over appropriate timescales for treatment of solid tumors.

CAR-T cells are most effective when fully activated at the tumor site^10, 11^, yet the high local cytokine concentrations required for CAR-T activation are highly toxic if delivered systemically, which typically precludes their use therapeutically. In current clinical approaches, T cells are expanded in high concentrations of cytokines only prior to administration into patients to avoid these toxicities. Locoregional approaches for delivering cytokines can limit systemic cytokine exposure and reduce toxicities^12^, but often involve engineering the cytokine itself through protein engineering approaches^13^. To date, the materials that have been designed for the co-delivery of T cells and cytokines have been limited to surgically implantable scaffolds, and the cytokines have had to be specially formulated either in microspheres or insoluble forms within the biomaterial to prevent rapid release and enable co-delivery of the therapeutic cells with these low molecular weight proteins^6, 8^.

Based on the promising results from previous studies of locoregional delivery of both CAR-T cells and cytokines, we hypothesized that next generation biomaterials affording enhanced control over the co-delivery of CAR-T cells and immunostimulatory molecules have the potential to transform solid tumor treatment strategies. For this approach we drew inspiration from recent reports in the field of tissue engineering, which suggest that locoregional delivery of stem cells in physically-crosslinked hydrogel scaffolds improves cell viability during injection, enhances cell retention at the targeted injection site and facilitates local expansion of transplanted cells^14–16^. In this work, we engineered a self-assembled and injectable biomaterials platform for CAR-T cell delivery based on Polymer-Nanoparticle (PNP) hydrogels^15, 17–21^ (Figure 1). These hydrogels leverage highly scalable chemistries^22^ and can be formulated under mild conditions that facilitate facile encapsulation of CAR-T cells and cytokines, while their shear-thinning and injectability enable minimally-invasive locoregional delivery of the therapeutic cells to tumors (Figure 1a). We showed that these injectable hydrogels can create a local inflammatory niche that prolongs CAR-T cell and cytokine exposure, thereby improving local CAR-T cell expansion and improving anti-tumor efficacy.

**Figure 1.**
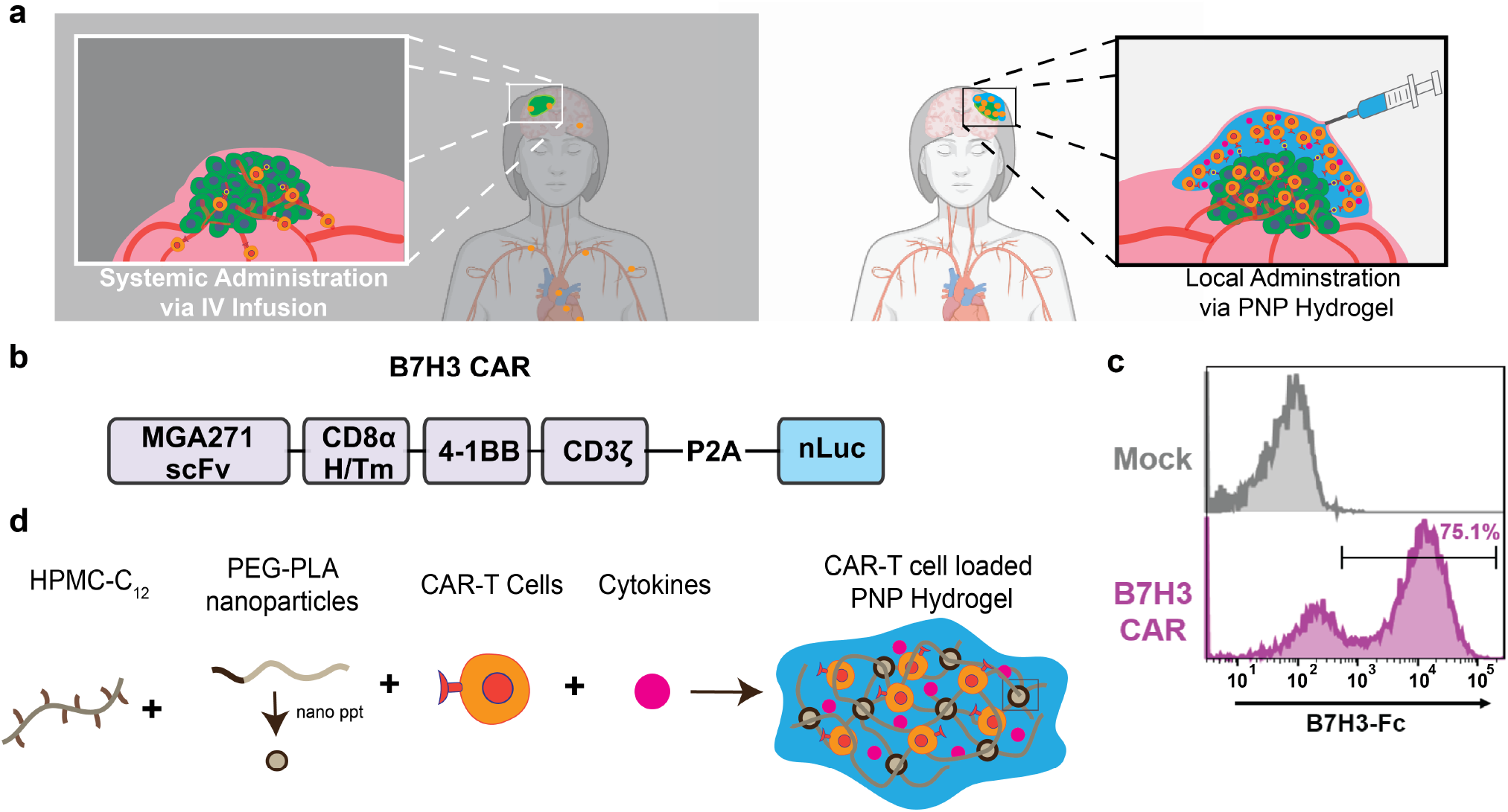
Injectable hydrogels for creation of a local inflammatory niche for co-delivery for CAR-T cells and cytokines to improve treatment of solid tumors. **a**, Schematic illustration demonstrating our proposed delivery method for CAR-T cells to solid tumors (right) compared to tradition intravenous approaches (left). **b**, Schematic illustration of the B7H3 CAR construct used for all studies. **c**, Transduction efficiency of B7H3 CAR-T Cells as determined by staining with B7H3-Fc compard to non-transduced “Mock” T cells. **d**, Formation of PNP hydrogels to co-encapsulate CAR-T cells and stimulatory cytokines through self-assembly of dodecyl-modified hydroxypropylmethylcellulose and degradable block-copolymer nanoparticles.

## Results and Discussion

### Development of an injectable hydrogel to act as an inflammatory niche

There are inherent materials design challenges in engineering hydrogels enabling both slow delivery of diverse immunostimulatory molecules while maintaining CAR-T cell viability. We have recently shown that the dynamic polymer mesh within PNP hydrogels can be engineered to be small enough to provide prolonged retention of proteins while still enabling immune cell motility^23^. We hypothesized, therefore, that co-encapsulation of immunostimulatory signals such as proliferative cytokines with CAR-T cells will enhance local T cell expansion following administration in the body.

To fabricate PNP hydrogels, a solution of dodecyl-modified hydroxypropylmethylcellulose (HPMC-C_12_) can be simply mixed with a solution of biodegradable nanoparticles comprising poly(ethylene glycol)-b-poly(lactic acid) (PEG-PLA NPs)^15, 17^. Upon mixing, dynamic multivalent and entropy-driven interactions between the HPMC-C_12_ polymers and the PEG-PLA NPs cause physical crosslinking and formation of a robust hydrogel material (Figure 2a-d). The self-assembled, entropy-driven crosslinking within these PNP hydrogels gives rise to temperature invariant mechanical properties^24^, and their physical crosslinking enables facile injection through a needle or catheter^25^. The mesh size of PNP hydrogels can be engineered to be sufficiently small to retain local immunomodulatory signalling over prolonged and controlled timescales^23^. To promote cellular motility and viability, the cell adhesion motif arginine-glycine-aspartic acid (RGD) can be attached to the hydrophilic corona of the PEG-PLA NPs using a “click” reaction prior to NP formation by nanoprecipitation of the PEG-PLA copolymers^15, 26^. We have previously shown that cells can be easily suspended in RGD-PEG-PLA NP solutions prior to mixing with HPMC-C_12_ for facile encapsulation of viable therapeutic cells^15^.

**Figure 2.**
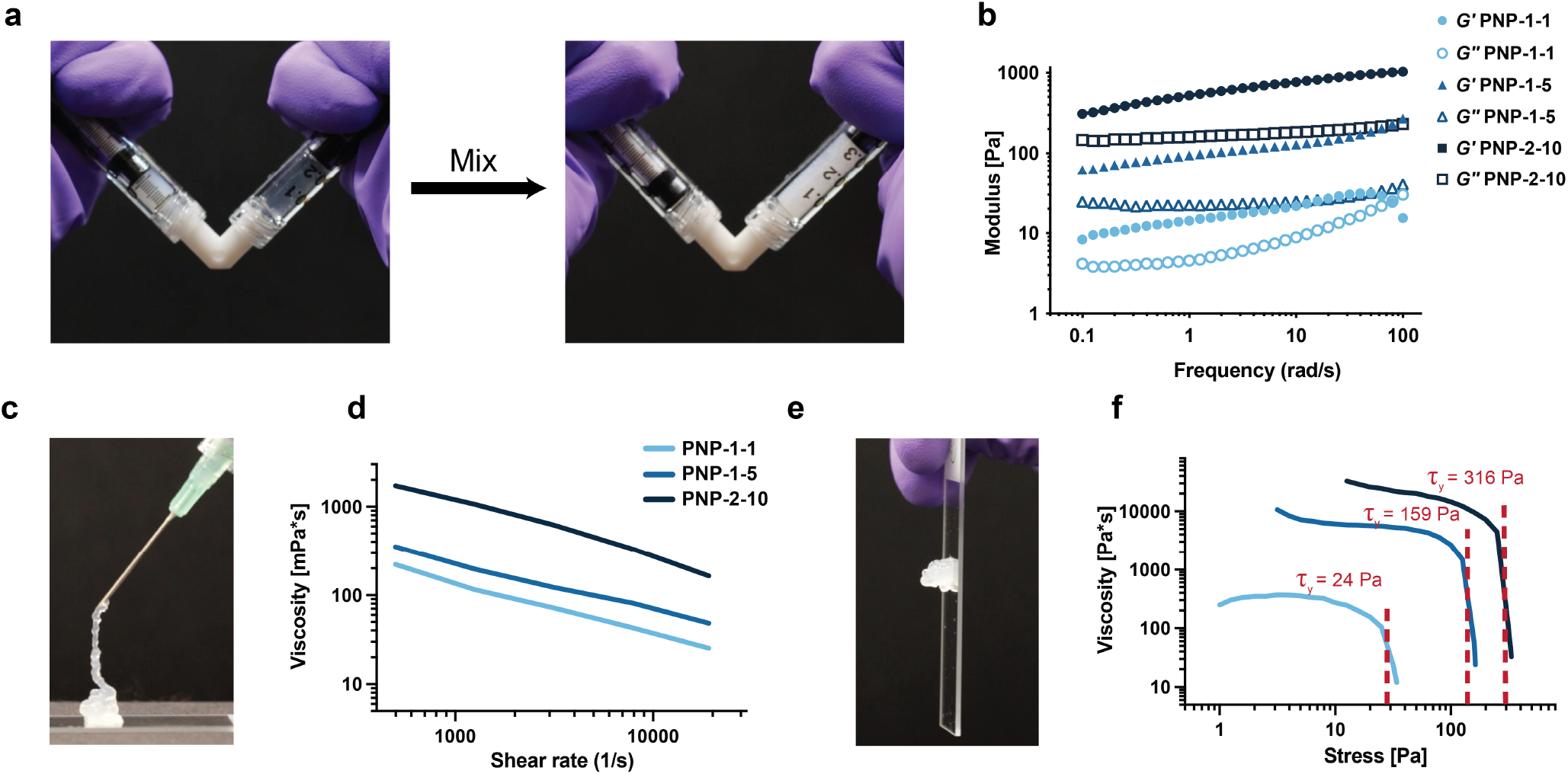
Rheological properties of PNP hydrogels enable injectability and depot formation. **a**, Facile formulation of PNP hydrogels by simple mixing of the biopolymer solution in one syringe (left) and the RGD-modified nanoparticles, cells and cytokines in the other syringe (right) using a luer-lock mixer. After gentle mixing for 30 seconds, a solid-like PNP hydrogel encapsulating cells homogeneously is formed (right syringe). **b**, Frequency sweep at 1% strain of three PNP hydrogels formulations, where first number is the wt% HPMC-C_12_ polymer and the second number is the wt% NPs (remaining mass as saline). **c**, Injection of cell-loaded PNP hydrogel through 26 G needle. **d**, Flow sweep at high shear rates (representative of injection) for three PNP formulations.**e**, A robust, solid-like hydrogel depot is formed that does not flow due to gravity. **f**, Stress sweeps for three PNP formulations demonstrating tunable yield stress flow behavior. Approximate yield stress transitions are noted with the red dotted lines.

We first sought to determine how hydrogel formulation affects the mechanical properties of PNP hydrogels (Figure 2e-h, Supplementary Figure 1). In this work we prepared three PNP hydrogel formulations referred to as PNP-1-1, PNP-1-5, and PNP-2-10, where the first number denotes the wt% of polymer and the second number denotes the wt% of nanoparticles (remaining mass is buffer). Oscillatory rheological testing demonstrated that these formulations all exhibit gel-like behavior across many timescales and up to high strains, as well as increasing stiffness with higher nanoparticle and polymer content (Figure 2e-f). Additionally, these formulations exhibited extreme shear-thinning without fracture at high shear rates that are representative of the injection process^25^. It is important to note that the shear-thinning behavior of this class of hydrogel not only enables injection through small diameter needles or catheters, but also protects encapsulated cells from harsh mechanical forces during the injection process, leading to higher cell viability upon delivery^27, 28^. After injection, the formulations rapidly self-heal to form robust depots with significant yield stress behavior that we hypothesized is critical for persisting in the subcutaneous space upon administration (Figure 2h).

### Prolonged retention of inflammatory cytokines in PNP hydrogels

It was first crucial that our materials platform exhibited degradation over relevant timescales for treatment of solid tumors. Previous described covalently-crosslinked hydrogel systems do not decay over relevant timescales and may persist long past treatment, increasing patient burden^6, 8^. To evaluate the persistence of our physically-crosslinked PNP hydrogel materials following implantation in mice, dye-labelled PNP-1-5 hydrogels were prepared by conjugation of a dye to the NPs used in fabrication of the materials, then implanted subcutaneously in mice by transcutaneous injection. The degradation of these materials was measured over time utilizing an *In Vivo* Imaging System (IVIS; Figure 3a-c). PNP-1-5 hydrogels degraded within weeks of implantation, exhibiting an average retention half-life of 8.9*±*2.6 days, which is consistent with the typical time-frame of a course of cancer treatment^29^.

**Figure 3.**
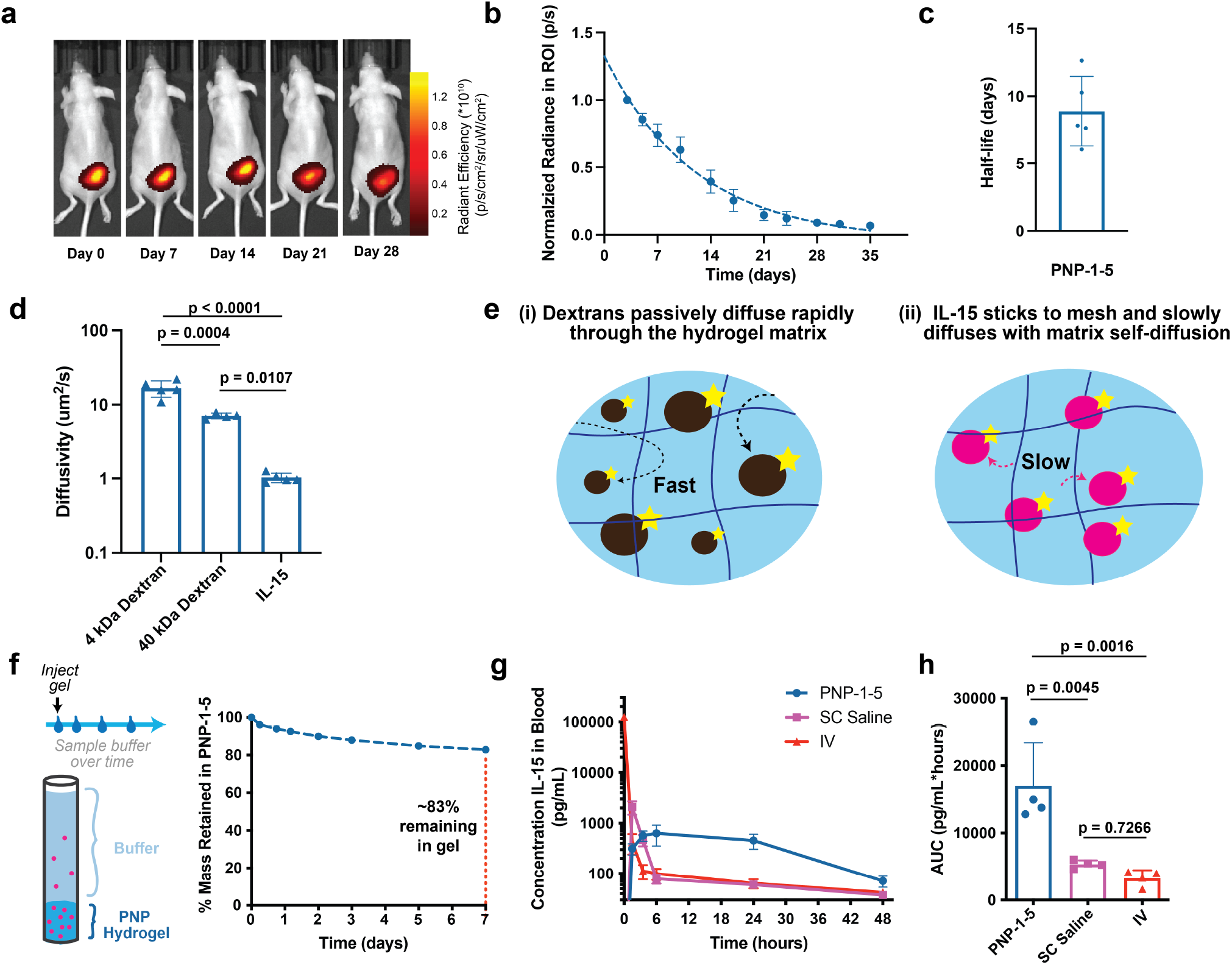
PNP hydrogels enable prolonged retention of stimulatory cytokines and exhibit controlled degradation *in vivo*. **a**, *In vivo* imaging of fluorescently-tagged PNP-1-5 hydrogel over 4 weeks after subcutaneous injection in mice. **b**, Quantification of fluorescence in the region of interest (ROI) surrounding the hydrogel and accompanying representative images over time. **c**, Degradation half-life of PNP-1-5 hydrogels *in vivo*. **d**, Diffusivity of FITC-dextran molecules and FITC-labelled IL-15 in the PNP-1-5 hydrogel formulation as measured by fluorescence recovery after photobleaching. **d**, Schematic illustrating the different diffusion mechanisms of the (i) dextran molecules and (ii) IL-15 in the PNP hydrogel mesh. **f**, Schematic of *in vitro* release assay of IL-15 from PNP-1-5 hydrogel immersed in saline over one week. % Mass of IL-15 remaining in the PNP-1-5 hydrogel during the release assay. **g**, Pharmacokinetic curves of IL-15 in the blood administered intravenously, subcutaneously, and from PNP-1-5 injected subcutaneously in mice. **h**, Area under the curve (AUC) of the pharmacokinetic profiles of the various IL-15 administration routes.

We hypothesized that co-encapsulation of CAR-T cells and stimulatory cytokines such as IL-15, a potent T cell activator that has also been found to support maintenance of T cell memory^30^, would improve CAR-T proliferation within the hydrogels. Moreover, as IL-15 is a small protein (15 kDa) that exhibits a short elimination half-life of only 1.5 hours in humans, we hypothesized that it would benefit from sustained retention to improve its exposure profile^30^. We therefore sought to determine whether PNP hydrogels could provide extended retention of IL-15 to improve CAR-T activation over longer timescales within the hydrogel following implantation. As mentioned above, cytokines can be easily encapsulated within these hydrogels by simple mixing during hydrogel fabrication without the need for complex formulation approaches or modification of the cytokine. We performed Fluorescence Recovery After Photobleaching (FRAP) experiments to assess diffusion of FITC-labelled IL-15 and FITC-labelled dextran polymers close in molecular weight to IL-15 (e.g., 4 kDa and 40 kDa). IL-15 exhibits a much lower diffusivity within PNP hydrogels than predicted for its low molecular weight and on the order of the self-diffusivity of the hydrogel mesh itself (Figure 3d)^20^, suggesting the IL-15 is non-specifically adhering to the hydrogel mesh (Figure 3d-e). This behavior likely arises from hydrophobic interactions between the IL-15 and the hydrophobically-modified cellulosic polymer mesh of the PNP hydrogels (Supplementary Figure 2a). *In vitro* release studies corroborated these observations, demonstrating that over 80% of the entrapped IL-15 was retained within the PNP-1-5 hydrogels after 1 week under infinite sink conditions (Figure 3f). While the IL-15 may interact with the hydrogel components, it was found to be highly stable and remained active in PNP hydrogels over prolonged timeframes under accelerated aging conditions (Supplementary Figure 2b). Furthermore, we conducted a pharmacokinetic study of IL-15 in mice following standard administration of the cytokine in a saline bolus or encapsulated in PNP-1-5 hydrogels. Intravenous (IV) and subcutaneous (SC) saline injections of IL-15 resulted in high maximum serum concentrations (C_*max*_), which is often associated with systemic toxicities, and rapid elimination. In contrast, sustained delivery of IL-15 from PNP-1-5 hydrogels significantly reduced the measured C_*max*_ values (C_*max,PNP*_ = 570*±*130 pg mL^-1^, C_*max,SC*_ = 2090*±*610 pg mL^-1^, p=0.0029), and enhanced total IL-15 exposure (AUC_*PNP*_ = 17000*±*6400 pg mL^-1^ h^-1^, AUC_*SC*_ = 5330*±*580 pg mL^-1^ h^-1^, p=0.0045) (Figure 3h-h).

### Development of an inflammatory niche for CAR-T cells

We next sought to determine how PNP hydrogel material properties affect CAR-T cell motility upon encapsulation (Figure 4a). Increasing the polymer and NP content within these hydrogels reduces the dynamic mesh size of the matrix and increases hydrogel stiffness^20^, both of which have been shown previously to impact the migration of T cells encapsulated within hydrogels^26^. While PNP hydrogels exhibit a small (on the order of nanometers) yet dynamic hydrogel mesh that inhibits passive diffusion of protein cargo, we hypothesized that this mesh would still enable active migration of encapsulated T cells^23^. To assess CAR-T cell motility within the PNP hydrogels, we conducted live cell motility experiments (Figure 4a-c). CAR-T cell migration speed decreased with decreasing hydrogel mesh size and increasing hydrogel stiffness (F_2,779_ = 97.47, p<0.001), suggesting migration and therefore retention within the hydrogel niche can be tuned through alteration of the hydrogel formulation (Figure 4b-e). We also found that RGD conjugation to the PEG-PLA NPs within the PNP hydrogel structure improves both T cell viability (p=0.0049) and mobility (p=0.009) (Supplementary Figure 2c-f).

**Figure 4.**
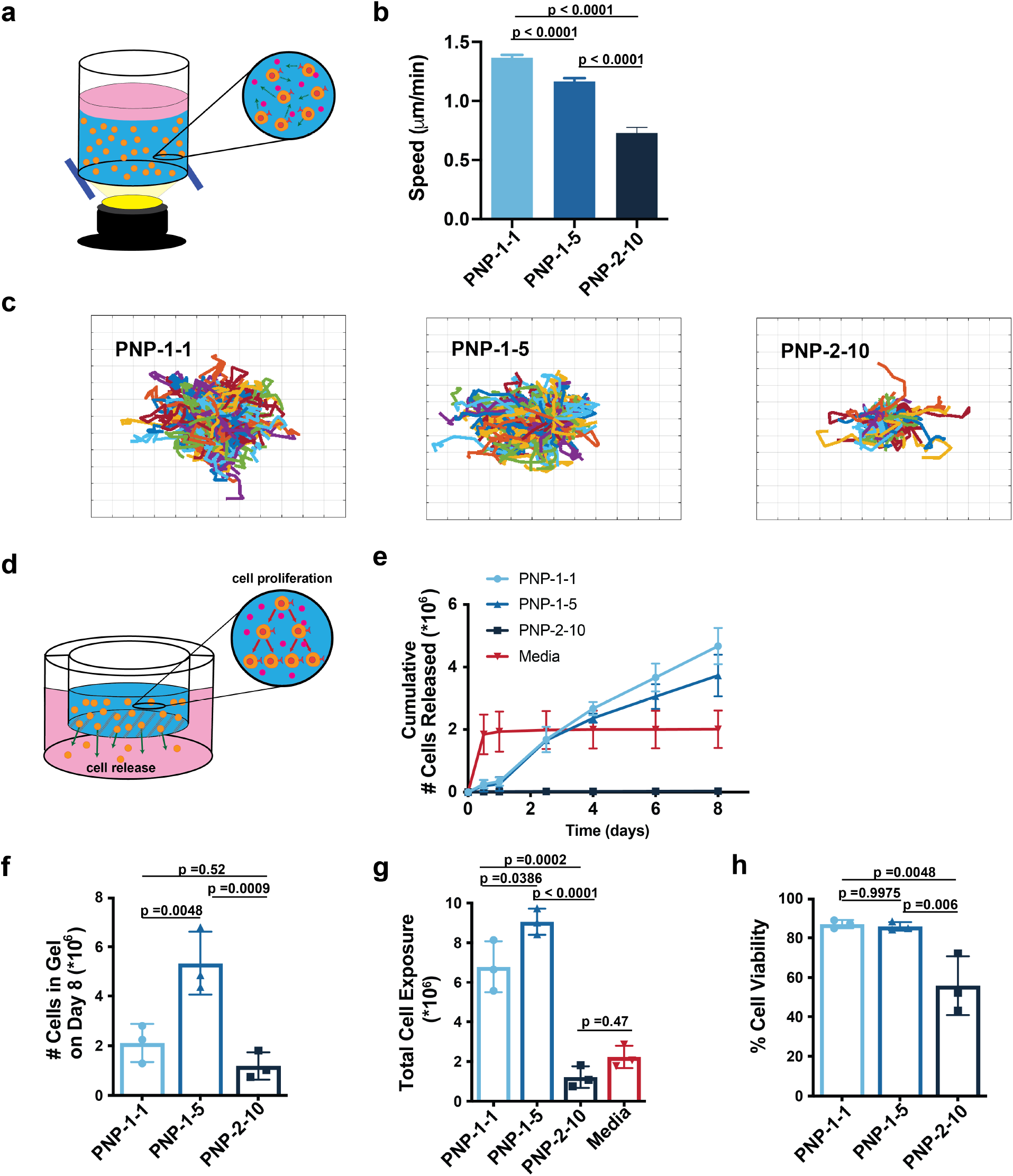
PNP hydrogels control T cell motility and release. **a**, Schematic illustration of *in vitro* experimental set-up to evaluate CAR-T cell motility within PNP hydrogels. **b**, CAR-T cell migration speeds in different PNP hydrogel formulations as quantified through cell migration experiments (n > 150 cells for all samples; mean *±* s.e.m.). **c**, Trajectories of migrating CAR-T cells within indicated formulations over 30 minutes. PNP-1-1, PNP-1-5, PNP-2-10 hydrogel formulations were tested, where first number is the wt% HPMC-C_12_ polymer and the second number is the wt% NPs. The trajectories are plotted at a common origin for easy visualization where each grid is 50 *μ*m wide. **f**, Schematic of experimental set-up to assess cell release from PNP hydrogel formulations. Hydrogel is injected into a porous transwell suspending over media. Cells both proliferate in the hydrogel and escape into the media below. **g**, Cumulative number of cells released into the media below over time of various PNP hydrogel formulations and a liquid media control. At each timepoint the media below the transwell was replaced. **h**, The total number of cells still remaining in the PNP hydrogel formulations at the end of the assay after 8 days, indicating if proliferation within the hydrogel in addition to release has occurred. **i**, The total cell exposure as quantified by the sum of the cumulative released cells and total cells in the transwell. **j**, Cell viability of the cells remaining in the PNP hydrogel formulations in the trans-well at the end of the cell release assay.

In contrast to tissue engineering applications in which cells are meant to remain within a scaffold, ACT requires that adoptive cells can effectively leave the depot. In addition to T cell motility within the hydrogels, we therefore also investigated migration of CAR-T cells out of the hydrogels. In these assays, CAR-T cells were loaded into various hydrogels and placed in trans-wells with a large pore size to allow T cells to escape the hydrogels and migrate into the media below (Figure 4d). The number of T cells that migrated into the media over time was determined using cell counting assays. We found that the PNP-2-10 hydrogel did not allow cellular egress over an 8-day period, while the PNP-1-1 and PNP-1-5 continuously released T cells over time (Figure 4e). In contrast, the bolus control released all of the T cells into the media within the first sampling period. At the end of the assay, when the hydrogels began to significantly degrade due to the immersion in media, hydrogels were dissociated and live cells still encapsulated within the gels were counted. The PNP-1-5 hydrogel formulation promoted the formation of an inflammatory niche that effectively expanded the CAR-T cells while enabling controlled migration of cells out of the hydrogel. In this way, the PNP-1-5 hydrogel enabled the highest total CAR-T cell exposure compared to the other hydrogel formulations and a 4.5-fold enhancement in total CAR-T cell exposure compared to standard bolus administration (p<0.0001; Figure 4f-h). For these reasons, the PNP-1-5 formulation was chosen to evaluate further in efficacy studies in a mouse cancer model.

### Evaluation of locoregional depot-based CAR-T therapy for solid tumors

To assess the ability of PNP hydrogels to improve CAR-T cell treatment of solid tumors *in vivo*, human B7H3 CAR-T cells (2 million/dose) were encapsulated in PNP-1-5 hydrogels with and without co-encapsulated IL-15 (0.25*/mu*g/dose) and administered peritumorally in NSG mice bearing a subcutaneous human medulloblastoma (MED8A) solid tumor^31, 32^ (Figure 5a, Supplementary Figures 3-4). Hydrogel-based treatments were compared to relevant controls, including IV and peritumoral adminstration of CAR-T cells with and without IL-15 (0.25 */mu*g/dose) in a saline vehicle. In these assays, both tumors and CAR-T cells were tracked and quantified in using a dual luciferase *in vivo* imaging approach using IVIS. As this model often exhibits severe Graft versus Host disease (GVHD) on account of the adoptive transfer of human T cells to NSG mice^33^, the length of the study is limited to 30 days.

**Figure 5.**
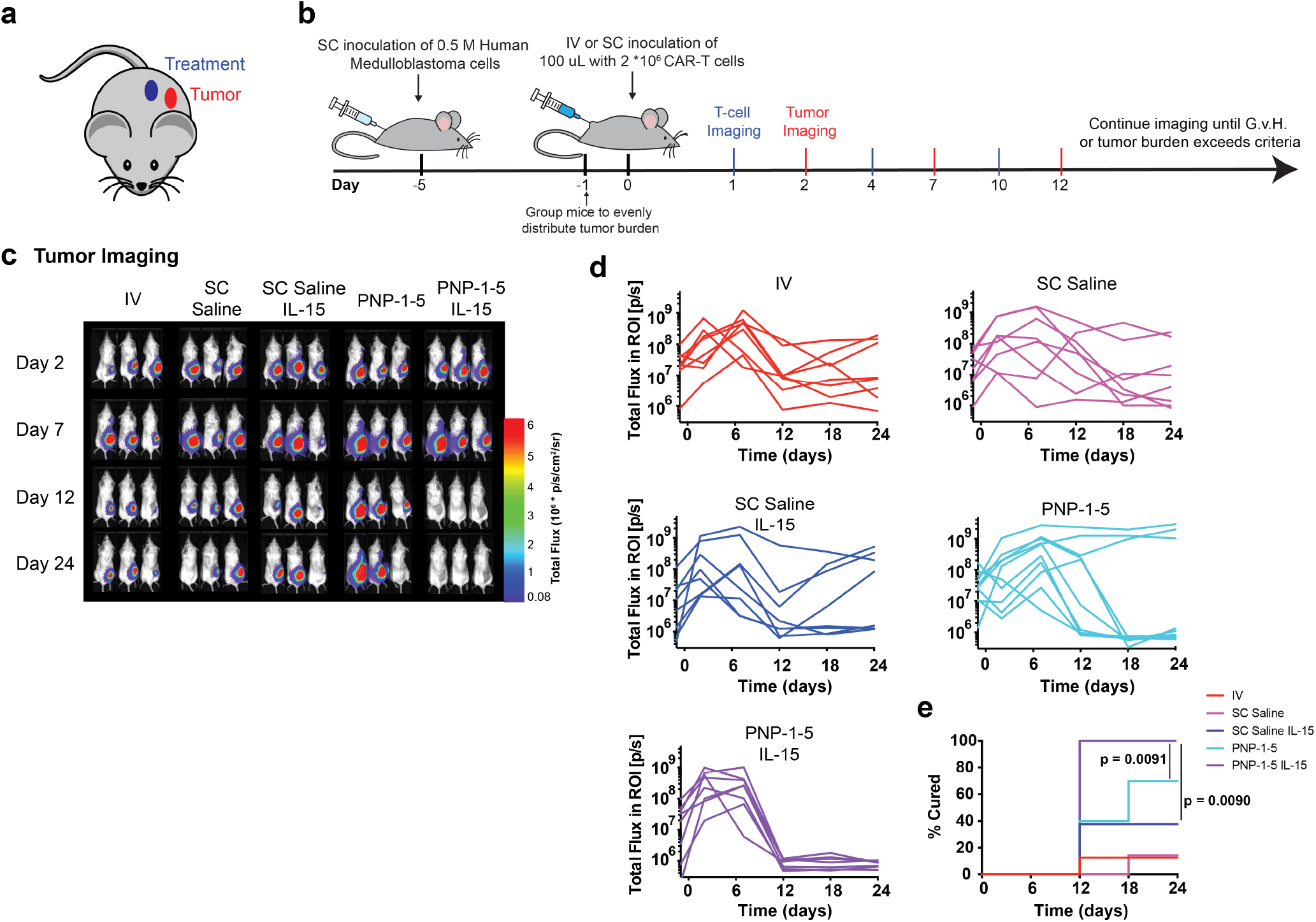
PNP Hydrogels improve treatment efficacy of solid tumors.. **a**, Schematic demonstrating placement of the tumor and the injected treatment. **b**, Experimental timeline for cancer experiments treating subcutaneous human medulloblastoma in mice with 2×10^6^ CAR-T cells administered with different delivery methods, including: (i) i.v. bolus, (ii) s.c. bolus, (iii) s.c. bolus delivery containing 0.25 *μ*g IL-15, (iv) PNP-1-5 hydrogel, and (v) PNP-1-5 hydrogel containing 0.25 *μ*g IL-15. **c**, Luminescent imaging of tumors in all experimental groups at all time points. **d**, Quantification of imaging data for all experimental groups (n=8-10 for all groups over 2 experiments). **e**, Percentage cured during the experiment across groups (defined as time when signal drops below and stays below 2*10^6^ p/s total flux). Further statistical analysis is reported in Supplementary Table 1.

While SC bolus and IV control treatments were only able to cure 10% and 40% of treated animals, PNP-1-5 hydrogel based treatments cured 70% of treated animals (p=0.023 and p=0.023, respectively; Supplementary Table 1). Indeed, even 4-fold higher CAR-T cell dosing adminstered SC or IV, demonstrated the inability to cure and cured at a slower rate than PNP-1-5 hydrogel delivery, suggesting a dose-sparing effect where PNP-1-5 hydrogels carrying fewer cells outperformed these more traditional approaches to CAR-T administration (Supplementary Figure 4).

Incorporation of IL-15 (0.25 *μ*g/mouse) within the PNP-1-5 hydrogels alongside the CAR-T cells further improved treatment (Figure 5c-d). The dose of IL-15 used in these studies was below the maximum tolerated subcutaneous dose of rhIL-15 used in recent human clinical trials scaled to mice (Supplementary Discussion)^12, 34, 35^, in contrast to many recent preclinical studies that utilize 5-20-fold higher concentrations of IL-15^8, 36^. Similarly, increased treatment efficacy was observed with co-delivery of IL-2 (0.25 *μ*g) alongside CAR-T cells in PNP-1-5 hydrogels (Supplementary Figure 5). It is also important to note that locoregional IL-15 delivery in PNP-1-5 hydrogels coupled with IV adminstration of “mock” CAR-T cells has no effect on tumor growth, confirming that CAR functionality is necessary for treatment and the locoregional exposure of IL-15 at this doses is insufficient to produce anti-tumor responses. Overall, co-delivery of CAR-T cells with stimulatory cytokines in PNP-1-5 hydrogels improved the efficacy and consistency of treatment, with all mice being completely cured by day 12 (Figure 5c-e). Indeed, while all mice treated with CAR-T cells and IL-15 in PNP-1-5 hydrogels were rapidly cured within the first two weeks, only 40% of mice receiving an SC bolus administration of cells and cytokines were cured by day 24 (p=0.009; Figure 5e). Interestingly, the efficacy of SC bolus administration of CAR-T cells was not significantly improved with co-delivery of IL-15 (p=0.29), further highlighting the benefit of generation of a local inflammatory niche within our PNP-1-5 hydrogels providing prolonged retention of IL-15 and CAR-T cells.

Enhanced T cell expansion was also observed when CAR-T cells were delivered in the PNP-1-5 hydrogels, particularly when co-delivered with IL-15 (Figure 6a-b). The PNP-1-5 formulation comprising IL-15 resulted in over a 100-fold increase in T cell expansion compared to the IV control (p=0.0417). Imaging data demonstrated that between days 10 and 21, the primary location of T cell signal moved from the hydrogel depot on the right subcutaneous flank to the spleen, aligning with the time-frame of tumor eradication. Histological analysis of the hydrogel depot confirmed that CAR-T cells are still present within the hydrogels several days after administration and demonstrated that no adverse immune response to the PNP hydrogel biomaterials were observable, consistent with previous findings^23^ (Supplementary Figure 6). Furthermore, analysis of inflammatory cytokines within the serum following treatment confirmed that locoregional hydrogel depot treatment did not elicit spikes in inflammatory cytokines that are often associated with systemic toxicities (Supplementary Figure 7), despite the greatly enhanced anti-tumor responses.

**Figure 6.**
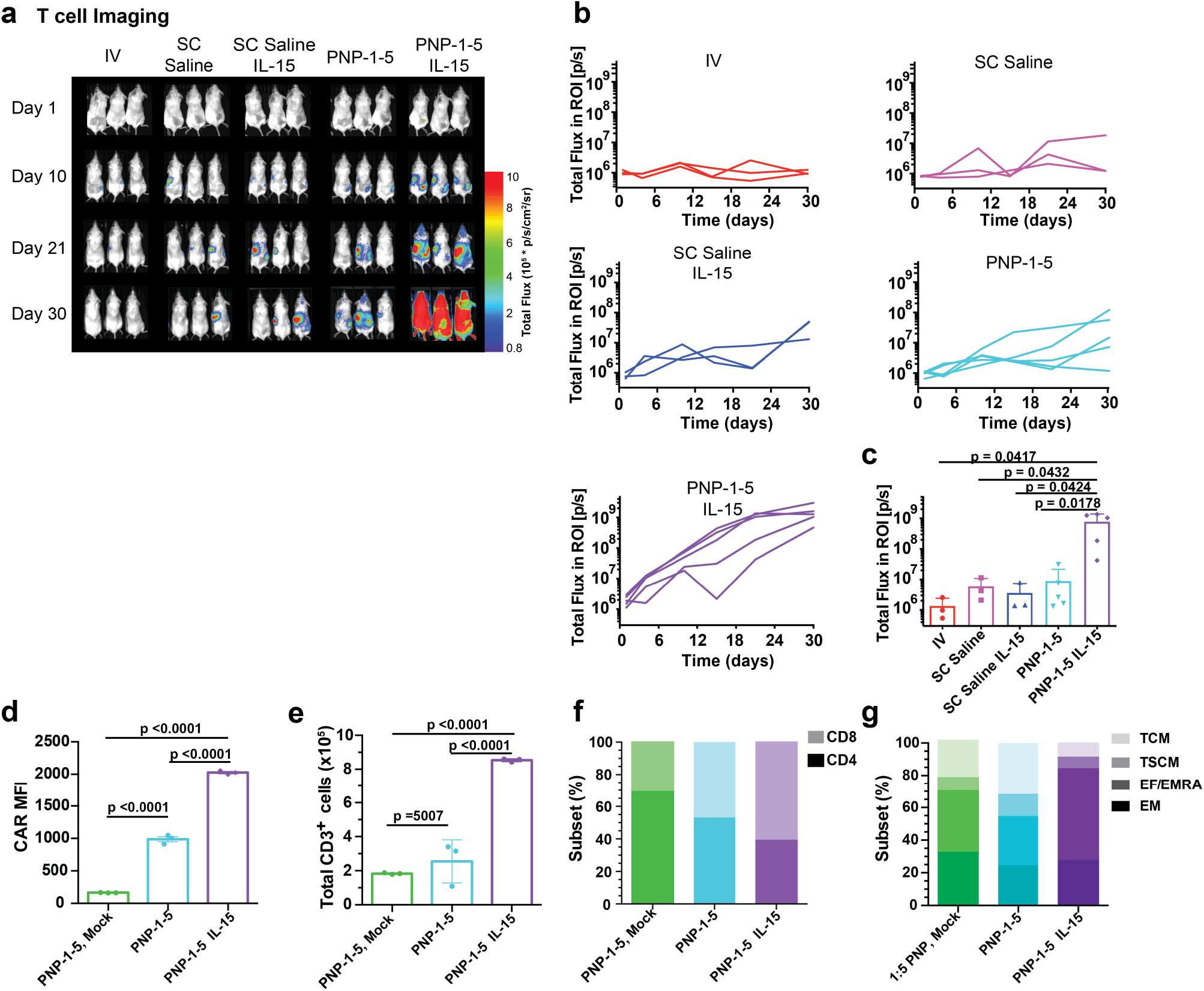
PNP Hydrogels improve expansion and phenotype of T cells. **a**, Luminescent imaging of CAR-T cells in all corresponding experimental groups. **b**, Quantification of imaging data for all experimental groups (n=3-5 for all groups). **c**, T cell signal across groups at Day 21. Additional P values for comparisons between all treatment groups can be found in Supplementary Table 2. **d**, CAR MFI (n=3 for all groups; mean *±* s.d.), **e**, total CD3+ T cells (n=3 replicates for all groups; mean *±* s.d.), **f**, relative CD4+ and CD8+ content (mean of n=3 for all groups), and **g**, memory CAR-T cell subsets (mean of n=3 replicates for all groups) of *ex vivo* expanded CAR-T cells extracted from PNP-1-5 gels 10 days after treatment in the MED8A tumor model.

**Figure 7.**
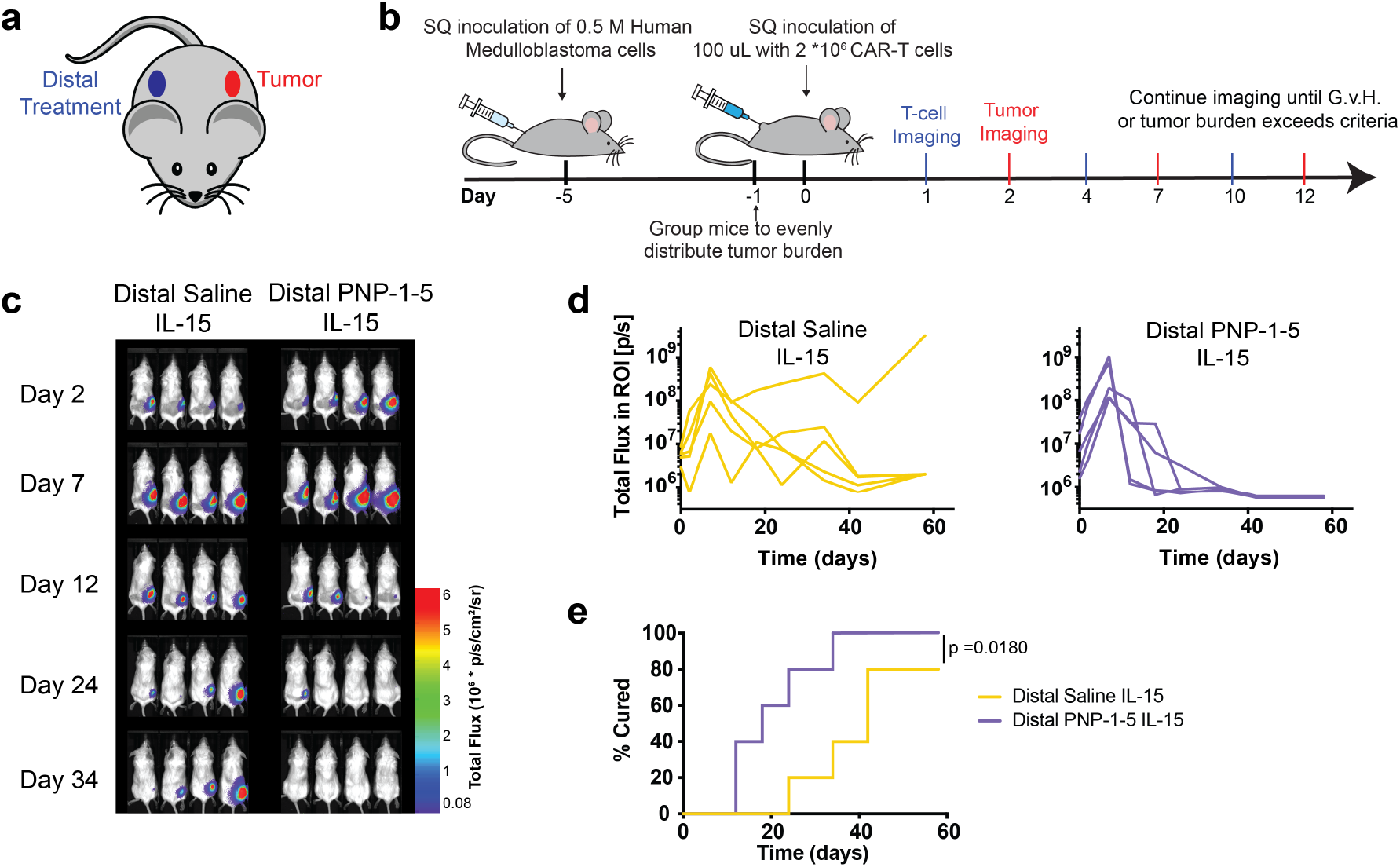
PNP Hydrogels are effective in treating distal subcutaneous human medulloblastoma in mice. **a**, Schematic showing placement of subcutaneous tumor and distal subcutaneous treatments. **b**, Schematic illustration of experimental timeline and treatment placement whereby mice received either (i) distal s.c. bolus injection of 2×10^6^ CAR-T cells and 0.25 *μ*g IL-15, or (ii) PNP-1-5 hydrogel containing 2×10^6^ CAR-T cells and 0.25 *μ*g IL-15. **c**, Representative luminescent imaging of tumors with *in vivo* imaging (n=5). **d**, Quantification of imaging data for both experimental groups (n=5). **e**, Percentage cured during the experiment across groups (defined as time when signal drops below and stays below 2*10^6^ p/s total flux).

### Evaluation of the impact of the local inflammatory niche on CAR-T phenotype

To better understand the mechanism for enhanced efficacy observed for co-delivery of CAR-T cells and IL-15 in a local inflammatory niche within PNP-1-5 gels, *ex vivo* analysis of T cells explanted from gels was performed 10 days after treatment. This analysis revealed enhanced CAR expression (p<0.0001) (Figure 6d), increased presence (p<0.0001) (Figure 6e), and greater proportions of CD8+ (Figure 6f) and T stem cell memory (TSCM) subsets (Figure 6f) when CAR-T cells were co-encapsulated with IL-15. Interestingly, non-significant changes to the proportions of central memory (TCM), effector memory (EM), and terminally-differentiated effector memory cells (EF/EMRA) T cell subsets were observed when IL-15 was co-encapsulated with CAR-T cells. Further, while all CAR-T treatment groups evaluated expressed similar levels of the activation markers 4-1BB, LAG-3, and PD-1, co-delivery of IL-15 within the PNP-1-5 hydrogels resulted in enhanced expression of CD39, which has recently emerged as an important marker of tumor-reactive T cells (Supplementary Figure 8)^37^. All memory subsets were also found to be similar between all CAR-T groups evaluated (Supplementary Figure 9).

### Evaluation of distal depot-based CAR-T therapy for solid tumors

In addition to delivering hydrogel-based CAR-T treatments peritumorally, we also sought to evaluate whether the hydrogel-based inflammatory niche can exhibit potent anti-tumor efficacy when delivered subcutaneously on the distal flank of the mouse from the tumor (Figure 7, Supplementary Figure 10). Distal administration ensures that CAR-T cells released from the hydrogel depot over time cannot drain directly to the tumor or the tumor-draining lymph node. We hypothesized that the enhanced expansion and maintenance of anti-tumor activation of the CAR-T cells enabled by the transient inflammatory niche within the hydrogels might elicit a potent abscopal effect relevant to metastatic cancer applications. *In vivo* imaging studies revealed that the hydrogel depot led to improved tumor clearance over a distal bolus control even though the two treatments exhibited similar overall T cell expansion and a similar distribution of T cell subsets in the blood and spleen (Supplementary Figure 11). Mice in these studies were monitored over a longer duration than the studies described above as they experienced significantly delayed GVHD. While all mice treated distally with CAR-T cells and IL-15 in PNP-1-5 hydrogels were rapidly cured within the first month, only 80% of mice receiving a distal SC bolus administration of cells and cytokines were cured by day 60 (p=0.018; Figure 7e). Overall, these results are promising as distal treatment with CAR-T cells and IL-15 within a transient inflammatory niche yields a potent abscopal effect that ultimately cures all treated animals, albeit slower than when this same treatment is given peritumorally. These results suggest that this hydrogel-based administration of CAR-T cells could have future use in treating metastatic cancers or inaccessible solid tumors.

### Outlook

While CAR-T therapy has resulted in remarkable anti-tumor effects in blood cancers, clinical successes in solid tumors have been severely limited as these tumors often exhibit an unfavorable microenvironment that can inhibit T cell infiltration and/or promote T cell dysfunction^1^. In this work we describe a CAR-T delivery method leveraging an injectable PNP hydrogel depot technology to generate a local inflammatory niche to expand and maintain a reservoir of CAR-T cells and stimulatory cytokines to improve their anti-tumor efficacy. The facile assembly and administration of these materials using standard injection approaches shows great promise for locoregional delivery of the CAR-T therapy. Cell motility and cytokine release studies demonstrated that these PNP hydrogels afford a unique opportunity to simultaneously inhibit the passive diffusion of cytokines while enabling active motility of encapsulated CAR-T cells, enabling formation of a transient inflammatory niche upon injection that enhances expansion of CAR-T cells in the body and induced a more tumor-reactive CAR-T phenotype compared to standard administration approaches. The *in vivo* expansion of more tumor-reactive CAR-T cells can potentially enable a reduction in the number of cells required for effective treatment, thereby reducing the high costs associated with the extended manufacturing periods currently required to generate sufficient cell numbers for standard treatment approaches. Moreover, the success of distal treatment of tumors with this transient inflammatory niche hold promise for development of treatments for inaccessible or metastatic solid tumors. Furthermore, this modular and versatile hydrogel depot technology is compatible with many cell and cytokine types as encapsulation and sustained delivery does not require alteration of either the cells or the cytokines, potentially enabling the use of these materials as the basis for treatments for many cancers. In future studies we aim to investigate formulations comprising other immunostimulants and checkpoint inhibitors to improve recruitment of endogenous immune cells to further amplify and expand anti-tumor responses. Overall, this interesting class of hydrogels addresses an unmet need for effective CAR-T cell delivery approaches that can enable the effective treatment of solid tumors.

## Methods

### Materials

All chemicals, reagents, and solvents were purchased as reagent grade from Sigma-Aldrich, Acros, or Alfa Aesar and used as received unless otherwise specified. Glassware and stir bars were oven-dried at 180 ^*°*^C. When specified, solvents were degassed by three cycles of freeze, pump, and thaw. HPMC-C_12_, PEG-PLA, and RGD-PEG-PLA were synthesized and characterized as described previously^15^. NPs were prepared by nanoprecipitation according to literature procedures using a 50:50 mixture of RGD-PEG-PLA:PEG-PLA polymers, and NP size and dispersity were characterized by dynamic light scattering (diameter = 35 nm, PDI = 0.02).^15^

### HPMC-C_12_ synthesis

Procedure was Hypromellose (HPMC) (1.5 g) was dissolved in N-methylpyrrolidone (NMP; 60 ml) by stirring at 80 ^*°*^C C for 1 h. Once the polymer had completely dissolved, the solution was cooled to room temperature. A solution of 1-dodecylisocyanate (0.5 mmol) was dissolved in NMP (5 ml) and added to the reaction mixture followed by 150 uL of NN-diisopropylethylamine as a catalyst. The solution was then stirred at room temperature for 16 h. This solution was then precipitated from acetone and the hydrophobically-modified HPMPC polymer was recovered by filtration, dried under vacuum at room temperature for 24 h and weighed, yielding HPMC-C_12_ as a white amorphous powder.

### PEG-PLA synthesis

Procedure was followed and analyzed as described previously.^15^ PEG (0.25 g, 4.1 mmol) and DBU (10.6 mg, 10 ml, 1.0 mol% relative to LA) were dissolved in dichloromethane (DCM; 1.0 ml). LA (1.0 g, 6.9 mmol) was dissolved in DCM (3.5 ml) with mild heating. The LA solution was then added rapidly to the PEG/DBU solution and was allowed to stir rapidly for 10 min. The PEG-b-PLA copolymer was then recovered from the reaction medium by precipitation from excess 50:50 mixture cold diethyl ether and hexanes, collected by filtration, and dried under vacuum to yield a white amorphous polymer. DMF GPC: Mn (Ð) = 21 kDa (1.07).

### RGD-PEG-PLA synthesis

Procedure was followed and analyzed as described previously.^15^ A 20 mL scintillation vial was charged with propargyl-functional cGGGRGDSP (22.7 mg, 26.6 *μ*mol), azido-functional PEG-PLA (0.530 mg, 21.2 *μ*mol), and NMP (4 mL). The reaction mixture was sparged with nitrogen for 10 min, then a degassed solution (0.1 mL) containing CuBr (3.7 mg/mL) and THPTA (16 mg/mL) was added and the reaction mixture was sparged with nitrogen for a further 10 min. The reaction mixture was incubated for 16h at room temperature, then the polymer was precipitated from an excess of cold diethyl ether in a 50 mL centrifuge tube and recovered by centrifugation. The polymer was then dissolved in acetone and precipitated again into diethyl ether, recovered by filtration, and dried in vacuum.

### Nanoprecipitation

Procedure was followed and analyzed as described previously^15^. A solution (1 mL) of PEG-PLA in acetonitrile (50 mg/ml) was added dropwise to water (10 mL) at a stir rate of 600 rpm. NPs were purified by ultracentrifugation over a filter (molecular weight cut-off of 10 kDa; Millipore Amicon Ultra-15) followed by resuspension in water to a final concentration of 250 mg/ml. NP size and dispersity were characterized by DLS (diameter = 35 nm, PD = 0.02). A 50:50 mixture of RGD-functionalized PEG-PLA polymer (RGD-PEG-PLA) to unmodified PEG-PLA polymer was used to create the RGD-NPs.

### Hydrogel formulation and cell encapsulation

Procedure was followed and analyzed as described previously^15^. HPMC-C_12_ was dissolved in phosphate-buffered saline at 6 wt% and loaded into a 1 mL luer-lock syringe. A cell pellet containing the number of cells to reach the desired concentration in the final hydrogels was suspended in phosphate-buffered saline. A 20 wt% nanoparticle solution in PBS was then added to the cell suspension. A 30:70 mixture of RGD to plain NPs were used to form the hydrogel for studies containing RGD, yielding a 0.5 mM concentration of conjugated RGD in the gel. The CAR-T cell/nanoparticle solution was loaded into a 1 mL luer-lock syringe. The cell/nanoparticle syringe was then connected to a female-female mixing elbow and the solution was moved into the elbow until it was visible through the other end of the elbow. The syringe containing the HPMC-C_12_ polymer was then attached to the elbow other end of the elbow. The two solutions were then mixed gently back and forth through the elbow for 30 seconds to 1 minute until the solutions had completely mixed and formed a homogeneous cell-loaded PNP hydrogel. IL-15 (R&D Systems) was incorporated with the nanoparticles during hydrogel formulation.

### Rheological characterization of hydrogels

Rheological testing was performed using a 20 mm diameter serrated parallel plate at a 600 *μ*m gap on a stress-controlled TA Instruments DHR-2 rheometer. All experiments were performed at 25^*°*^C. Frequency sweeps were performed at a strain of 1%. Amplitude sweeps were performed at frequency of 10 rad/s. Flow sweeps were performed from low to high stress and steady state sensing.

### Viscometry at high shear rates

A Rheosense m-VROC viscometer was used to measure the hydrogel viscosity at high shear rates from low to high using a 1 mL Hamilton syringe. Each data point was collected at steady state.

### Cell lines

MED8A was kindly provided by S. Chesier (Stanford University, Stanford, CA). MED8A-GFP-Fluc cells were cultured in DMEM supplemented with 20% FBS, 100 U/mL penicillin, 100 *μ*g/mL streptomycin, 2 mM L-glutamine, and 10 mM HEPES (Gibco). STR DNA profiling was conducted once per year (Genetica Cell Line testing) and routinely tested for mycoplasma. Cell lines were cultured in a 5% CO_2_ environment at 37^*°*^C.

### Plasmid construction and virus production

B7H3 CAR-P2A-Nluc plasmid was constructed by fusing the MGA271 scFv to CD8*α* hinge and transmembrane, 4-1BB costimulation domain, CD3*ζ* signaling domain, porcine teschovirus-1 2A (P2A) ribosomal skipping sequence, and nanoluc in an MSGV retroviral vector^38–40^. The Antares-P2A-mNG constructed was constructed by fusing a P2A sequence and mNeonGreen to the c-terminus of Antares^41, 42^. Retroviral supernatant was produced using 293GP packaging cells transfected with the RD114 envelope plasmid and the corresponding plasmid construct, as previously described^43^.

### CAR-T cell isolation

T cells were isolated from buffy coats purchased from the Stanford Blood Center under an IRB-exempt-protocol. Negative selection using the RosetteSep Human T cell Enrichment kit (Stem Cell Technologies) and SepMate-50 tubes was performed to purify primary human T cells. T cells were crysperserved in CryoStor CS10 media at a concentration of 1-2×10^7^ cells/mL.

### CAR-T manufacturing

Primary human T cells were thawed at Day 0 and activated with anti-CD3/CD28 Dynabeads (Thermo Fisher) at a 3:1 bead to T cell ratio and cultured in AIM V + 5 % heat-inactivated FBS, 100 U/mL penicillin, 100 mg/mL streptomycin, 2 mM L-glutamine, 10 mM HEPES, and 100 U/mL rhIL-2). On Day 2, virus-coated wells were prepared on 12-well non-tc, Retronectin-coated (Takara/Clonetech) plates by spinning 1 mL of the corresponding virus on at 3200 RPM for 2 hours. T cells were then cultured on these plates for 24 hours. This transduction process was on Day 3 after. Beads were magnetically removed on day 4, and cells were expanded until Day 10 for *in vivo* experiments and Day 10-14 for *in vitro* experiments. For dual virus cotransductions, T cells were transduced with Antares-P2A-mNG on Day 2 and B7H3 CAR-P2A-Nluc on Day 3.

### *Ex vivo* CAR-T cell analysis

For *ex vivo* analysis of transferred T cells, mice were euthanized ten days after T cell administration. Gels were harvested and T cells were extracted by mechanical dissociation (gentleMACS dissociator, Miltenyi). Single-cell suspensions were filtered and stained for flow cytometry.

### Flow cytometry

B7H3 CAR was detected using recombinant B7H3-Fc (R&D Systems) fluorescently labeled with the DyLight 650 Microscale Antibody Labeling Kit (Thermo Fisher). The following antibodies were used to stain T cells: BUV395 Mouse Anti-Human CD4 (Clone SK3, BD), BUV805 Mouse Anti-Human CD8 (Clone SK1, BD), BV605 Mouse Anti-Human CD62L (Clone DREG-56, BD), and BV711 Mouse Anti-Human CD45RA (Clone HI100, BD). CAR-T cell quantification was performed using the CountBright Absolute Counting Beads (Thermo Fisher). Flow cytometry was performed on a BD Fortessa and analyzed on FlowJo version 10.7.1.

### PNP hydrogel *in vivo* degradation study

AF647 NPs were prepared using a combination of PEG–PLA (25 mg) and unconjugated azide-PEG–PLA (25 mg). A 1 ml solution of combined PEG–PLA and azide-PEG–PLA in DMSO (50 mg/ml) was added dropwise to 10 ml of water at room temperature under a high stir rate (600 rpm). NPs were purified by ultracentrifugation over a filter (molecular weight cut-off of 10 kDa; Millipore Amicon Ultra-15) followed by resuspension in water to a final concentration of 200 mg/ml. The nanoparticles were then functionalized by mixing adize-functional NPs (250 *μ*L, 20 wt%) with AF647-DBCO (25 *μ*l, 1 mg/ml) and waiting 12 hours. PNP hydrogels were made according to procedures above using a 1:5 formulation. Each 100 *μ*L gel contained 5 l of fluorescent AF647-conjugated NPs and 20 *μ*L of non-fluorescently conjugated nanoparticles. Five SKH1E mice were each injected with 100 *μ*L of fluorescently tagged 1:5 PNP hydrogel and imaged using the in-vivo Imaging System (IVIS Lago) over a series of timepoints spanning 35 days. When imaged, mice were anesthetized with isoflurane gas and imaged with an exposure time of 0.25 seconds, excitation wavelength of 600 nm, and emission wavelength of 670 nm (binning: medium, F/stop: 1.2). Total radiant efficiency ([photons/s]/[μW/cm^2^]) was quantified using an equal-sized region of interest surrounding the gel depot. As early time points (time < 3 days) showed fluorescence in the region of interest to increase instead of decrease, fluorescent intensity at each timepoint was normalized to fluorescent intensity on day 3. Normalized fluorescence intensity values for each mouse (n=5) between day 3 and 35 were fit to a single exponential decay models and half-lives were acquired and averaged using GraphPad Prism.

### CAR-T cell proliferation study

Promega CellTiter-Glo 3D Cell Viability Assay was used to characterize the short-term cell proliferation in different formulation conditions. Cells were encapsulated at 1 million cells per 100 *μ*L in an opaque 96 well plate. Relative signal was measured after 2 days in culture by adding 100 *μ*L per well of the CellTiter-Glo reagent, mixing for 5 minutes, allowing the plate to sit for 25 minutes, and then reading the luminescent signal with a 1 second integration time. Relative growth was calculated using a calibration curve generated by testing various known cell loadings in hydrogel.

### CAR-T cell migration studies

Live cell imaging was done with a laser scanning confocal microscope (Leica SP8) fitted with temperature/incubator control, suitable for live-imaging (37^*°*^C, 5% CO_2_). A 20x-air objective, NA = 0.75, was used to acquire approximately 60*μ*m stack images for 5 hours (at 10 minute intervals). Imaging parameters were adjusted to minimize photo-bleaching and avoid cell death. After completion of migration studies, the centroids of mCherry-labeled cells were tracked using the spots detection functionality in Imaris (Bitplane). Poorly segmented cells and cell debris were excluded from the analysis and drift correction was implemented where appropriate. A custom MATLAB script was used to reconstruct cell migration trajectory.

### CAR-T cell hydrogel release studies

CAR-T cells were loaded into PNP hydrogel formulations at 20 million cells/mL and 100 *μ*L hydrogel was injected into transwell inserts. Transwells were gently placed into 24 well plate, and 650 *μ*L media was gently added below. The experiment was incubated according to standard culture conditions. At each timepoint, the media was gently removed, the cells within the media were counted, and the transwell was placed in a new well with new media. At the end of the assay (8 days, when hydrogel was beginning to noticeably dissociate) the contents of the transwell were diluted, cells were counted, and viability was recorded.

### Fluorescence recovery after photobleaching

Fluorescein Isothiocyanate-Dextran (4 kDa and 40 kDa, Sigma) were encapsulated at 1 mg/mL in PNP hydrogel formulations. IL-15 cytokine was labelled with using a FITC Conjugation Kit (Fast)-Lightning-Link kit (abcam) and encapsulated in PNP hydrogels at 8 *μ*g/mL. Gels were placed onto glass slides and imaged using a confocal LSM780 microscope. Samples were imaged using a low intensity laser to observe an initial level of fluorescence. Then the laser was switched to full intensity and focused on a region of interest (ROI) with a 25 *μ*m diameter for 10 seconds in order to bleach a circular area. Fluorescence data were then recorded for 4 minutes to create an exponential fluorescence recovery curve. Samples were taken from different regions of each gel (n=5-9). The diffusion coefficient was calculated according to the following equation:

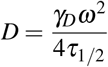

where the constant 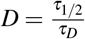 with *τ*_1/2_ being the half-time of the recovery, *τ*_*D*_ the characteristic diffusion time, both yielded by the ZEN software, and *ω* the radius of the bleached ROI (12.5 *μ*m)^23, 44^.

### *In vivo* experimental approaches

All experiments followed protocols approved by the Stanford Administrative Panel on Laboratory Animal Care. 5×10^5^ MED8A-GFP-Fluc cells were injected s.c. on the right flank of NSG mice 5 days prior to treatment in a 50:50 ratio of Matrigel (Cultrex Pathclear) to phosphate buffered saline (PBS). One day prior to treatment mice were imaged and distributed into groups of roughly equivalent tumor burden. CAR-T cells were encapsulated in phosphate buffered saline or PNP hydrogels in 1 mL syringes in a tissue culture hood one hour before injection as previously described. One syringe was prepared for each experimental group within a study. 300 *μ*L extra gel was prepared in each syringe. Double the volume of bolus and intravenous controls was prepared. Syringes were transported on ice to the animal facility. 100 *μ*L of phosphate buffered saline or PNP hydrogel was delivered to each mouse. Subcutaneous injections were delivered in a 21 G luer lock syringe.

### *In vivo* imaging

To image tumors, mice were intraperitoneally injected with D-Luciferin, potassium salt (Goldbio) at 150 mg/kg in phosphate buffered saline. After 5 minutes, mice were anesthetized with isoflurane gas and imaged with an exposure time of 30 seconds with an In Vivo Imaging System (Spectral Imaging Instruments Lago-X). Signal was quantified as the total flux of photons/sec in the region of interest at peak intensity. The region of interest was defined as a rectangular box of consistent size around the entire mouse. Background signal for quantification was defined as the maximum signal observed through all imaging experiments in an equivalently sized rectangular box with no luminescent signal. To image tumors, mice were intraperitoneally injected with nano-luciferin (NanoLuc, Promega) at a 40x dilution in phosphate buffered saline. After 5 minutes, mice were anesthetized with isoflurane gas and imaged with an exposure time of 30 seconds In Vivo Imaging System (Spectral Imaging Instruments Lago-X). Signal was quantified as the total flux of photons/sec in the region of interest at peak intensity. The region of interest was defined as a rectangular box of consistent size around the entire mouse.

### Histology

Gels were explanted through disection from mice on Day 5 of treatment and frozen in optimal cutting temperature compound (OCT). All samples were processed and stained by Stanford Animal Histology Services with hematoxylin and eosin staining and Cy5-CD3 staining. Two replicates were collected from PNP-1-5 hydrogel containing IL-15, and two replicates were collected for PNP-1-5 hydrogel. Representative images are shown.

### Cytokine release *in vitro*

Capillary tubes were incubated with 0.1 wt% Bovine serum albumin overnight at 5 ^*°*^C. Capillary tubes were then dried through using house air and evaporation and then loaded with 100 *μ*L of PNP hydrogel containing 0.25 *μ*g IL-15. 300 *μ*L phosphate buffered saline (PBS) was loaded on top of each gel. Samples were stored at 37 ^*°*^C to mimic physiological environments. At each time point, the PBS was completely removed using a long needle and stored at -80 ^*°*^C for later analysis. The PBS was then replaced. IL-15 concentrations were determined by ELISA according to the manufacturer’s instructions (R&D Systems Human IL-15 Quantikine Assay). Absorbance was measured at 450 nm in a Synergy H1 Microplate Reader (BioTek). At the end of 7 days, the gel was diluted by 10 in saline, allowed to incubate for 3 hours, and analyzed for the remaining cytokine. Cytokine concentrations were calculated from the standard curves. Mass in gel was calculated as the inverse of the total mass released into the release buffer during the study and the cytokine left in the gel at the end of the study.

### Cytokine stability *in vitro*

IL-15 was encapsulated in PNP-1-5 hydrogel at 2.5 *μ*g/mL. 100 *μ*L hydrogel was loaded in low-binding eppendorf tubes. Eppendorf tubes were incubated at 37 ^*°*^C and shaking at 150 rpm until various timepoints. At each timepoint tubes were removed from the incubator, diluted by 10 with saline containing 0.1 wt% BSA and 5 wt% trehalose and stored at -80 ^*°*^C until further analysis. IL-15 concentrations were later determined by ELISA according to the manufacturer’s instructions (R&D Systems Human IL-15 Quantikine Assay). Absorbance was measured at 450 nm in a Synergy H1 Microplate Reader (BioTek). Cytokine concentrations were calculated from the standard curves.

### Cytokine release *in vivo*

Serum was collected at the indicated times by tail vein blood collection and stored at -80 ^*°*^C. Serum IL-15 concentrations were determined by ELISA according to the manufacturer’s instructions (R&D Systems Human IL-15 Quantikine Assay). Absorbance was measured at 450 nm in a Synergy H1 Microplate Reader (BioTek). Cytokine concentrations were calculated from the standard curves.

### Inflammatory cytokine analysis

60 *μ*L blood was collected after 48 hours from each mouse (n=3 for each group). Samples were processed and analyzed by the Stanford Human Immune Monitoring Center. Samples are reported relative to naive mice.

### Statistical methods

All error is reported as standard deviation unless reported otherwise. Sample size for each experiment is included in the corresponding methods section as well as figure captions. Significance is reported with p values from a one-way ANOVA followed by Tukey’s post hoc test for multiple comparisons. Statistical analyses on percentage-cured plots were conducted using a log-rank Mantel-Cox test.

## Author’s contributions

A.K.G. and E.A.A. conceived of the idea. A.K.G. and L.L. performed experiments. Other authors aided with experiments. All authors contributed towards the manuscript.

## Acknowledgements

This research was financially supported by the Center for Human Systems Immunology with Bill and Melinda Gates Foundation (OPP1113682) as well as the American Cancer Society (RSG-18-133-01). A.K.G.is thankful for a National Science Foundation Graduate Research Fellowship and the Gabilan Fellowship of the Stanford Graduate Fellowship in Science and Engineering. L.L. was supported by a Stanford Graduate Fellowship, a National Science Foundation Graduate Research Fellowship (DGE-1656518), and a Siebel Scholar fellowship. S.C. was supported by the National Cancer Institute of the National Institutes of Health under Award Number F32CA247352. C.L.M. was supported by the NSERC Postgraduate Scholarship and the Stanford BioX Bowes Graduate Student Fellowship. E.C.G. was supported by NIH Cell and Molecular Biology Training Program (T32 GM007276). B.S.O. was supported by the Eastman Kodak Fellowship. A.I.D. was supported by the Schmidt Science Fellows program, in partnership with the Rhodes Trust. O.A. was supported by National Institutes of Health F31 (1F31CA250405-01). O.C. was supported by National Institutes of Health National Cancer Institute Grant (R37 CA214136). We would like to acknowledge the Stanford Center for Innovations in In vivo Imaging (small animal imaging center), particularly Jason Thanh Lee and Laura J. Pisani. We would also like to acknowledge the Stanford Animal Histology Core, particularly Eric Petersen.

