## Supplemental Information for "Delivery of CAR-T Cells in a Transient Injectable Stimulatory Hydrogel Niche Improves Treatment of Solid Tumors"

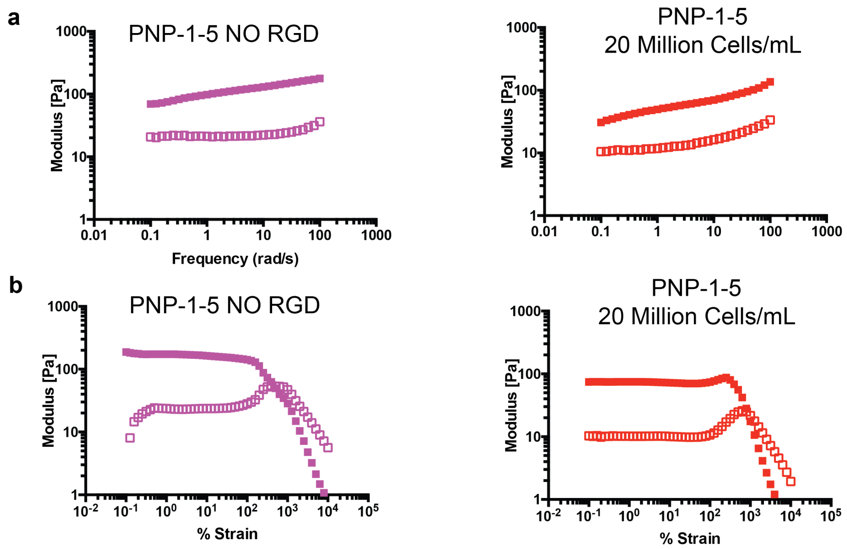


**Supplementary Figure 1:** Rheology of the PNP-1-5 hydrogel control formulations including controls of hydrogels without RGD-conjugated nanoparticles and hydrogels containing 20 million cells/mL. **a,** Frequency sweep (%strain=1%) for all formulations at 25°C. **b,** Amplitude sweep ($\omega$=10 rad/s) for all formulations at 25°C.

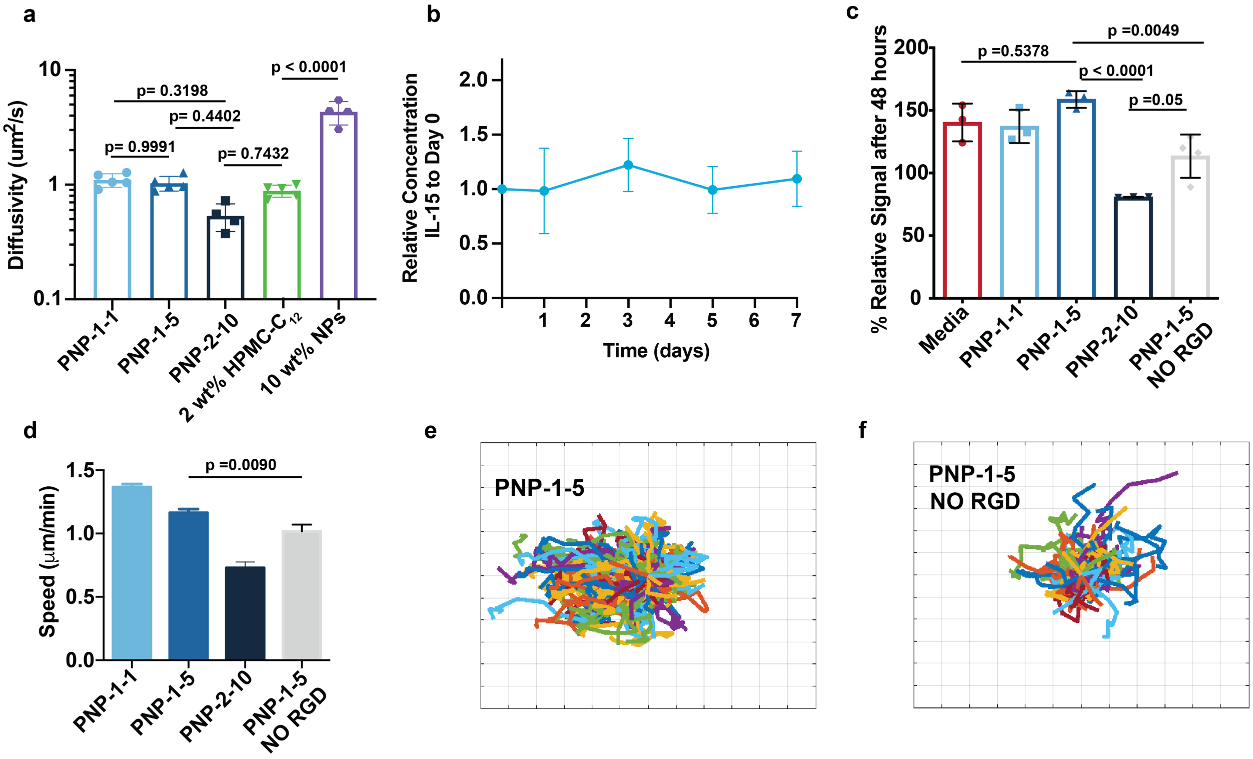


**Supplementary Figure 2:** *In vitro* experiments assessing diffusivity and stability IL-15 cytokine in PNP hydrogels and the benefits of RGD-conjugation for T cell proliferation and motility. **a,** Diffusivity of IL-15 in various PNP hydrogel formulations and components (first number is polymer wt%, second number is NP wt% (remaining mass is saline). **b,** Cytokine stability in PNP-1-5 hydrogel over time relative to time zero assessed by ELISA. **b,** Relative proliferation after 48 hours in PNP hydrogel formulations and 2D control. **c,** CAR-T cell speeds within PNP-1-5 hydrogel formulations with and without conjugation of RGD moieties (data shown as mean$\pm$SEM). **d/e,** Trajectories of migrating CAR-T cells within indicated hydrogel formulations. The trajectories are plotted at a common origin for easy visualization. Each grid represents 50 $\mu$m.

**Supplementary Table 1:** Additional P values from Figure 5e survival analysis computed using a log-rank Mantel-Cox test.

| Treatment pair | P value |
|---|---|
| IV, SC Saline | 0.9624 |
| IV, SC Saline + IL-15 | 0.2626 |
| IV, PNP-1-5 | 0.023 |
| IV, PNP-1-5 + IL-15 | 0.0006 |
| SC Saline, SC Saline + IL-15 | 0.2954 |
| SC Saline, PNP-1-5 | 0.023 |
| SC Saline, PNP-1-5 + IL-15 | 0.0002 |
| SC Saline + IL-15, PNP-1-5 | 0.1655 |

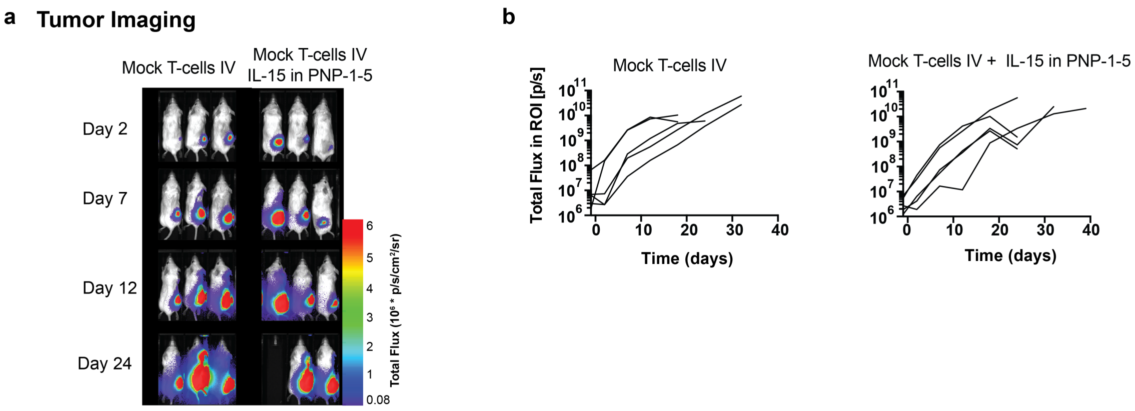


**Supplementary Figure 3:** Related *in vivo* controls confirming the CAR functionality is essential for treatment in the *in vivo* model. **a,** One group containing non-transduced "Mock" T cells (2 million) delivered intravenously, another group containing non-transduced "Mock" T cells (2 million) delivered intravenously with additional PNP-1-5 hydrogel with a 0.25 $\mu$g dose of IL-15 injected subcutaneously (gel not containing any cells). Tumor imaging using an *in vivo* imaging system. **b,** Corresponding quantification of luminescent signal from tumor imaging (n=5 for both groups from one experiment).

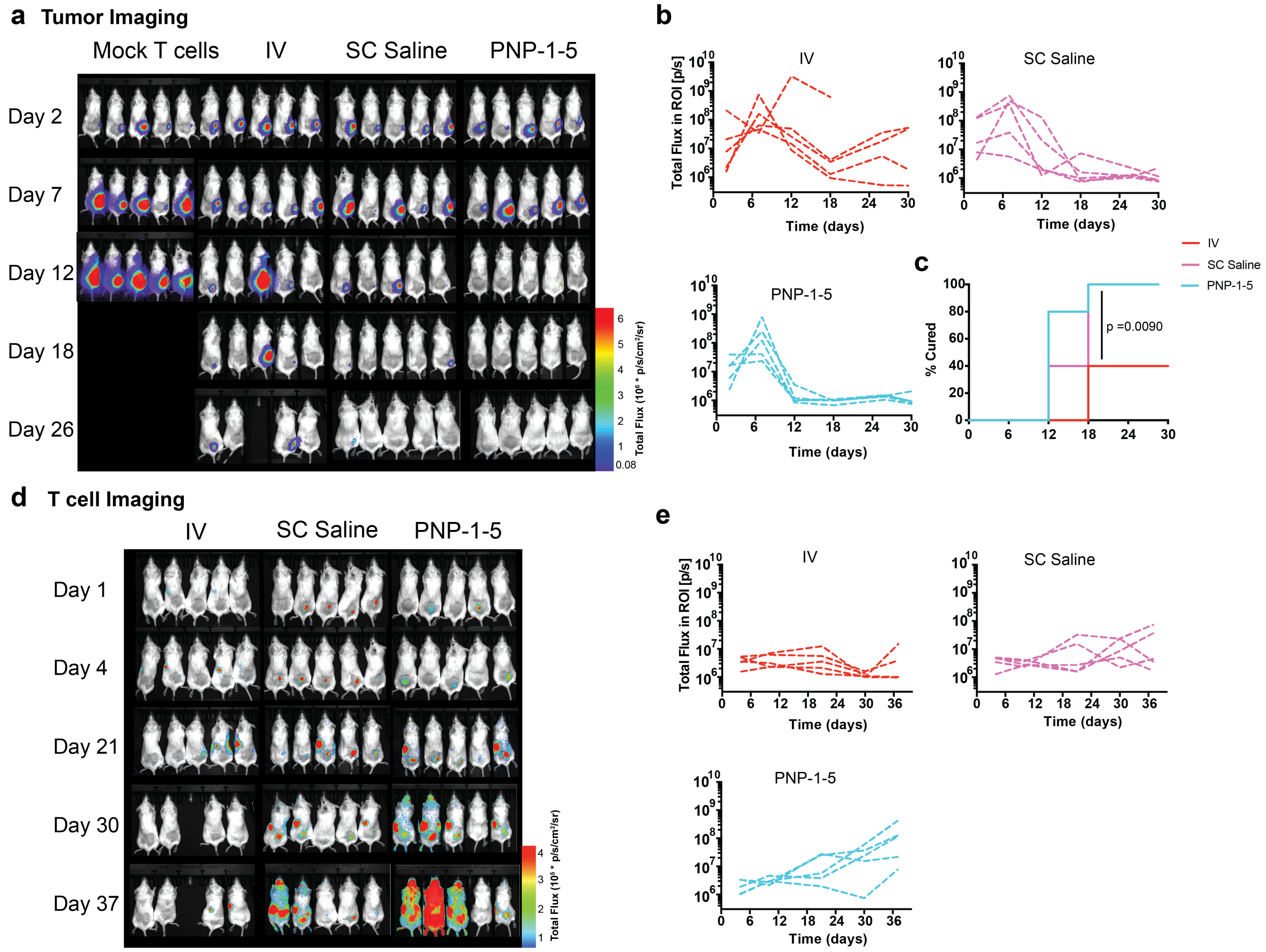


**Supplementary Figure 4:** *In vivo* experiment delivering 8 million CAR-T cells in PNP hydrogels and controls (cytokines not included). **a,** Tumor imaging using an *in vivo* imaging system (n=5 from one experiment for all groups). **b,** Corresponding quantification of luminescent signal from tumor imaging. **c,** Percentage cured during the experiment across groups (defined as time when signal drops below and stays below $2*10^6$ p/s total flux). P = 0.02 for IV compared to SC Saline. P = 0.22 for PNP-1-5 compared to SC Saline. **d,** CAR-T cell imaging using an *in vivo* imaging system. **e,** Corresponding quantification of luminescent signal from CAR-T cell imaging (n=5 from one experiment for all groups).

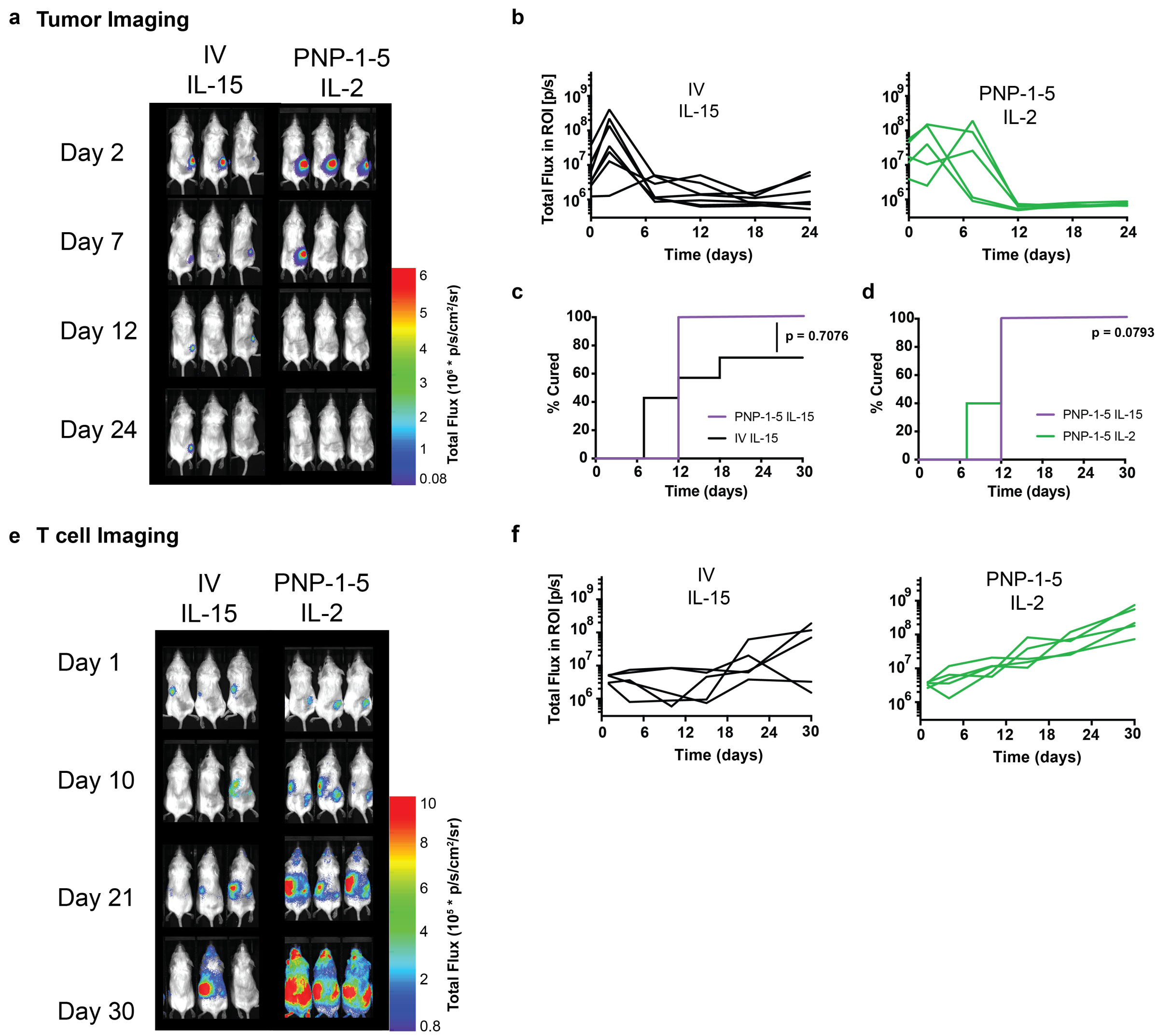


**Supplementary Figure 5:** Related *in vivo* controls confirming benefits to co-delivery of cells and cytokines and PNP hydrogels. This intravenous dose of IL-15 exceeds the human scaled maximum tolerated dose found in patients with cancer and would not be a translationally relevant treatment strategy.[1] **a,** CAR-T cells (2 million) co-delivered with IL-15 intravenously at a 0.25 $\mu$g dose of IL-15, and CAR-T cells (2 million) encapsulated in PNP-1-5 hydrogel with a 0.25 $\mu$g dose of IL-2. Tumor imaging using an *in vivo* imaging system. **b,** Corresponding quantification of luminescent signal from tumor imaging (n=7 from one experiment for IV group, n=5 for PNP-1-5 with IL-2 from one experiment). **c,** Percentage cured during the experiment in the IV IL-15 group compared the PNP-1-5 IL-15 group (defined as time when signal drops below and stays below $2*10^6$ p/s total flux). **d,** Percentage cured during the experiment in the PNP-1-5 IL-2 group compared to the PNP-1-5 IL-15 group (defined as time when signal drops below and stays below $2*10^6$ p/s total flux). **e,** CAR-T cell imaging using an *in vivo* imaging system (n=5 for all groups). **f,** Corresponding quantification of luminescent signal from CAR-T cell imaging (n=5 for both groups from one experiment).

**Supplementary Table 2:** Additional P values for treatment groups from Figure 6c calculated using a one-way ANOVA with a posthoc Tukey multiple comparisons test.

| Treatment pair | P value |
|---|---|
| IV, SC Saline | >0.9999 |
| IV, SC Saline + IL-15 | >0.9999 |
| IV, PNP-1-5 | >0.9999 |
| SC Saline, SC Saline + IL-15 | >0.9999 |
| SC Saline, PNP-1-5 | >0.9999 |
| SC Saline + IL-15, PNP-1-5 | >0.9999 |

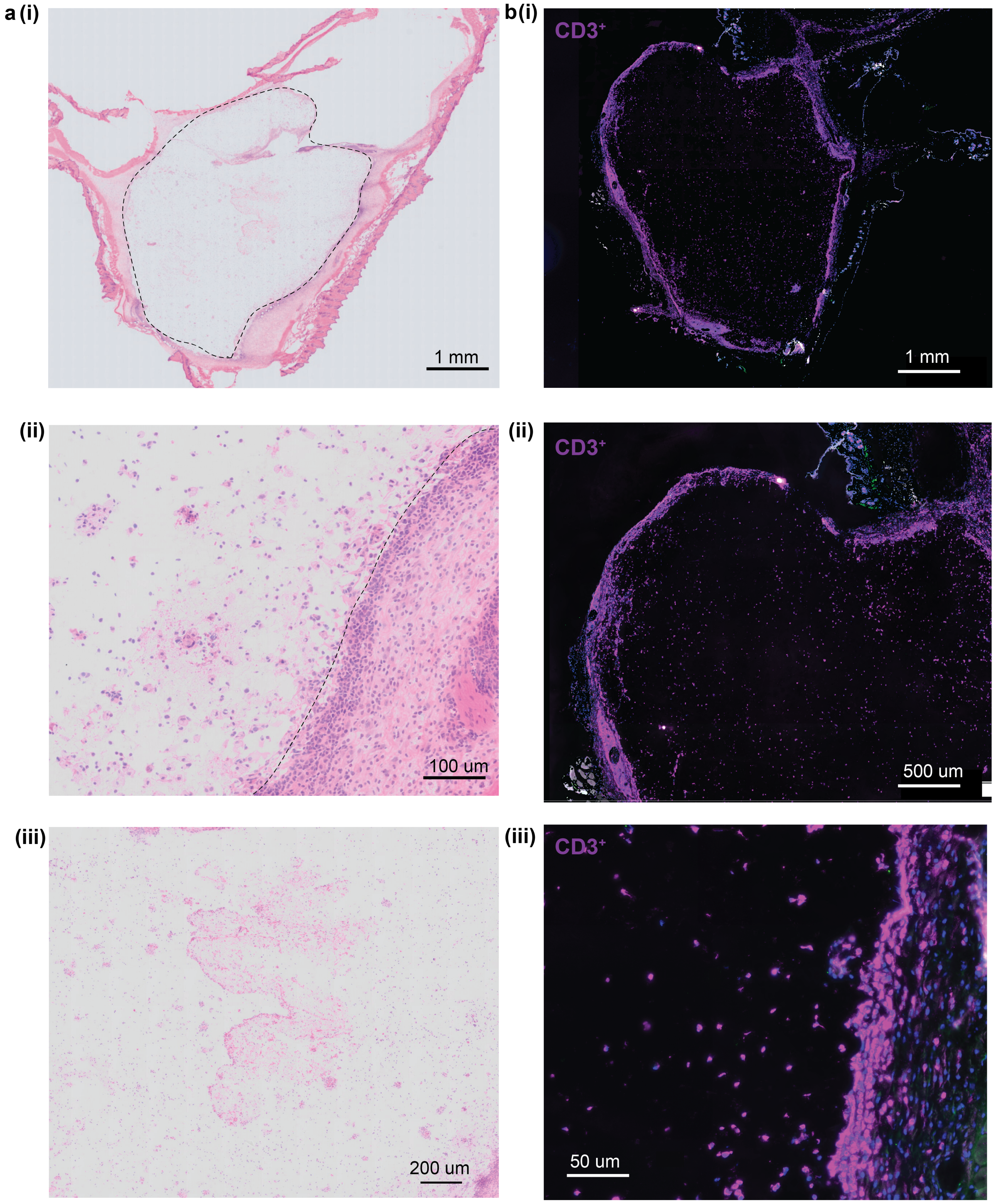


**Supplementary Figure 6:** Histology of explanted PNP-1-5 hydrogel containing CAR-T cells (2 million) and IL-15 after 5 days *in vivo*. **a,** Images of Hematoxylin and Eosin staining under various magnifications (indicated by scale bars). Hydrogel is outlined with black dotted lines and surrounded by exogenous tissue and cells. **b,** Images of $CD3^+$ staining in purple (Cy5, Jackson Immuno) under various magnifications (indicated by scale bars). $CD3^+$ staining includes recruited endogenous cells in addition to the delivered cells.

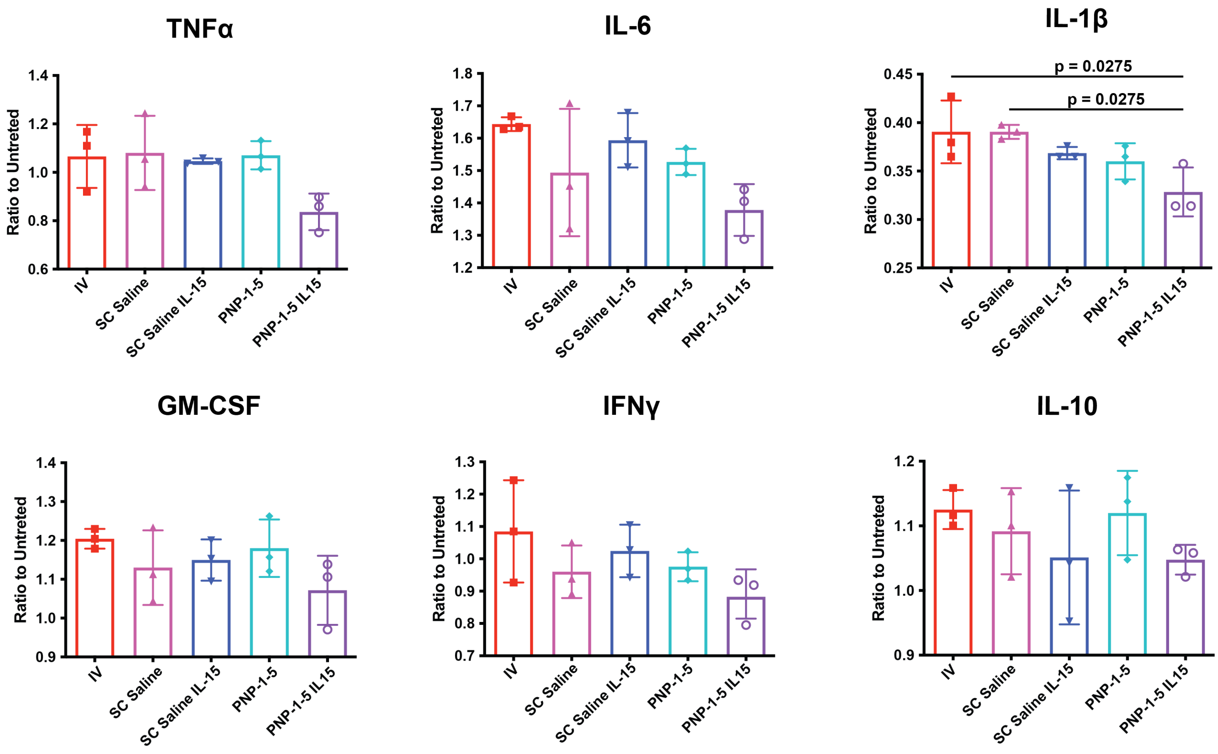


**Supplementary Figure 7:** Inflammatory mouse cytokines measured in the blood 3 days after treatment with CAR-T cells (2 million) in various delivery methods reported relative to values determined for naive mice (n=3 mice per group). P values are only reported here if $p<0.05$.

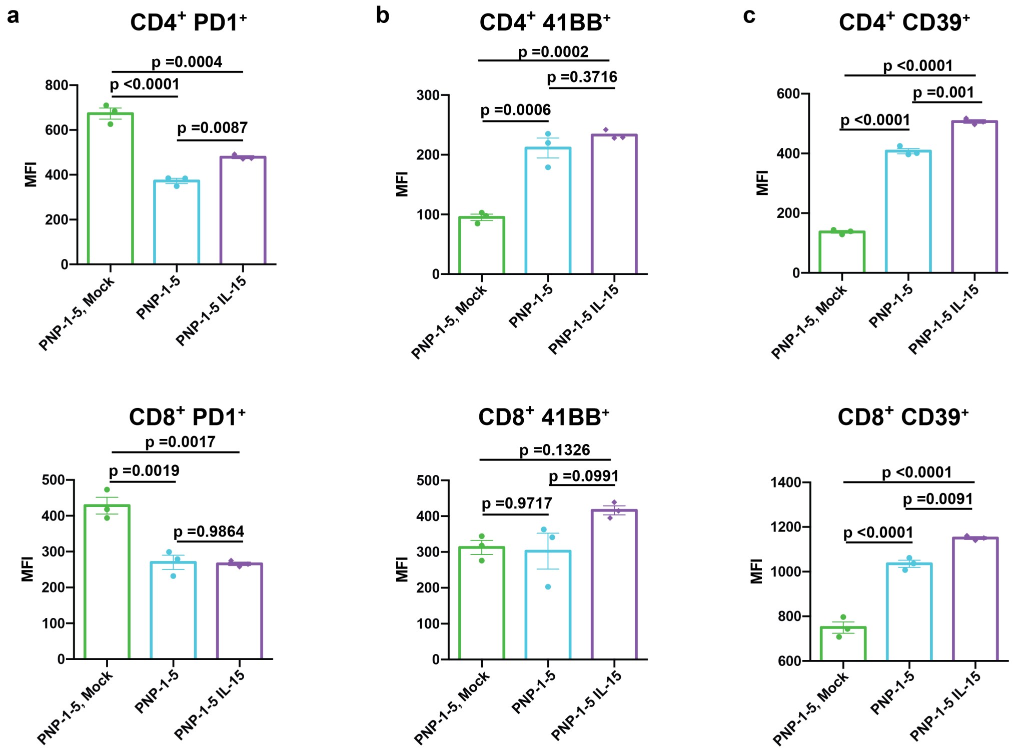


**Supplementary Figure 8:** Expression of T cell activation markers on CAR-T cells extracted from PNP hydrogels. MFI of **a,** PD1, **b,** 4-1BB, and **c,** CD39 staining on CAR-T cells collected 10 days after treatment in the MED8A tumor model. Data shown as mean±SEM (n=3).

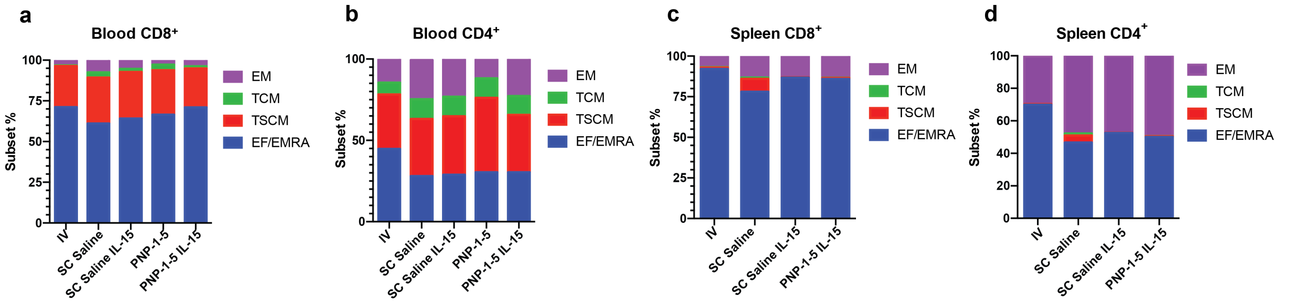


**Supplementary Figure 9:** T cell memory subsets from CAR-T cell treated mice. T cell memory subsets, as determined by CD62L and CD45RA staining, from **a, b,** blood and **c, d,** spleen samples collected 10 days after CAR-T cell administration in the MED8A tumor model.

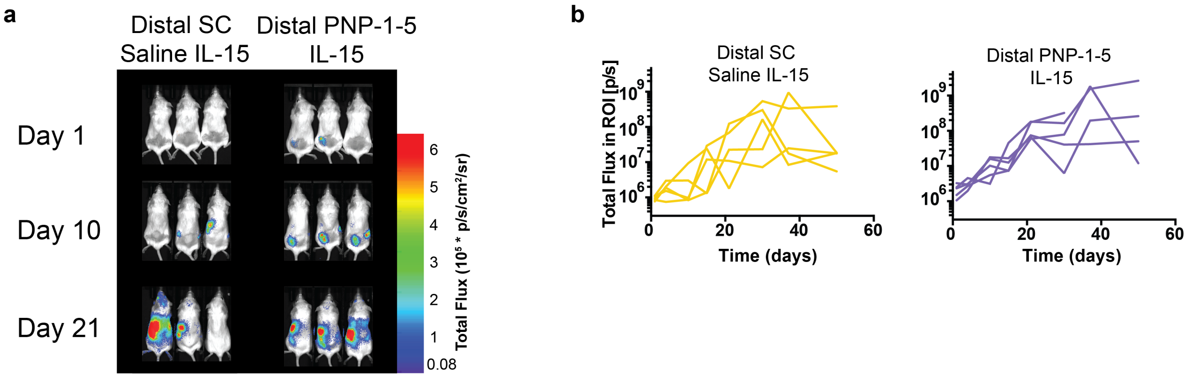


**Supplementary Figure 10:** *In vivo* experiment comparing CAR-T cell expansion when administered subcutaneously on the contralateral (left) flank, distal to the tumor (right subcutaneous flank). CAR-T cells (2 million) were co-administered in PNP-1-5 hydrogels or in a saline bolus at a dose of 0.25 $\mu$g/mouse IL-15. **a,** CAR-T cell imaging using an *in vivo* imaging system (n=5 for all groups). **b,** Corresponding quantification of luminescent signal from CAR-T cell imaging (n=5 for all groups).

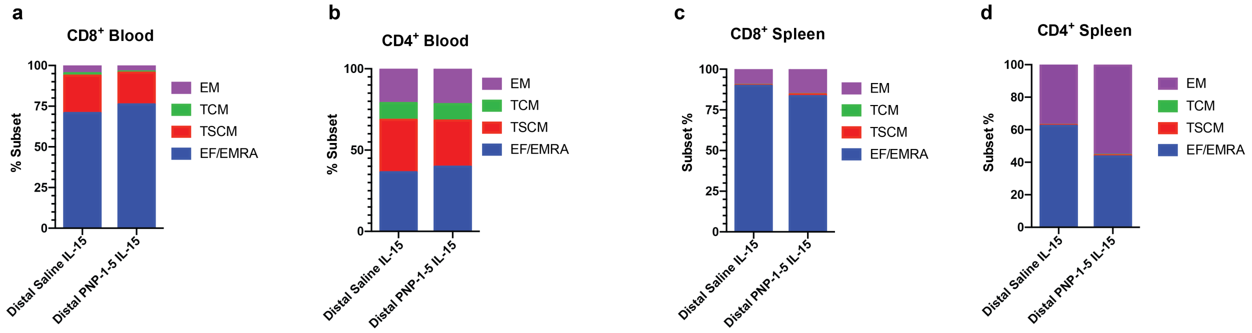


**Supplementary Figure 11:** CAR-T cell memory subsets from distally treated mice. CAR-T cell memory subsets, as determined by CD62L and CD45RA staining, from **a,b,** blood and **c,d,** spleen samples. CAR-T cells were collected 10 days after treatment in the MED8A tumor model. Data shown as mean$\pm$SEM (n=3).

### Supplementary Discussion

#### Calculation of rhIL-15 Dose Equivalence in Humans

While many CAR-T therapies require lymphodepletion, there has been little research on how lymphodepletion affects the maximum tolerated dose of rhIL-15, so the dosage in our studies is based on the available literature. The maximum tolerated subcutaneous dose per day in recent human clinical trials is 2 $\mu$g/kg/day.[2] We can scale this dose to mice by multiplying by 12.3, giving 24.6 $\mu$g/kg/day.[3] Assuming mice are approximately 0.02 kg in weight, gives 0.492 $\mu$g/day in one dose in a mouse. We chose to administer approximately half this dose, 0.25 $\mu$g, subcutaneously in our studies. Most preclinical studies to date deliver higher doses of IL-15.[4–6] Note that even the dose we use in our studies exceeds the maximum tolerated dose found in patients with cancer of rhIL-15 intravenously delivered in humans, 0.3 $\mu$g/kg/day.[1]
